# Characterizing cytosine methylation of polymorphic human transposable element insertions using human pangenome resources

**DOI:** 10.1101/2024.11.22.623804

**Authors:** Xiaoyu Zhuo, Chad Tomlinson, Edward A. Belter, Prashant Kumar Kuntala, Wesley N. Saintilnord, Juan Jiang, Tina Lindsay, Juan Macias, Human Pangenome Reference Consortium, Robert S. Fulton, Ting Wang

**Affiliations:** Department of Genetics, Washington University School of Medicine, St. Louis, MO, USA; The Edison Family Center for Genome Sciences and Systems Biology, Washington University School of Medicine, St. Louis, MO, USA; McDonnell Genome Institute, Washington University School of Medicine, St. Louis, MO, USA

## Abstract

Cytosine methylation, a crucial epigenetic modification, plays a vital role in genomic regulation. Leveraging the advancements in third-generation sequencing, we investigated the methylation patterns of non-reference insertions of human lymphoblastoid cell lines (LCLs), particularly polymorphic transposable elements (TEs). We validated the high concordance between long-read methylation calls and conventional whole genome bisulfite sequencing (WGBS) method. By characterizing thousands of polymorphic TE insertions genome-wide using long reads from the draft Human Pangenome Reference, we aimed to establish general rules of TE methylation by addressing three key questions: 1) what is the methylation profile of each insertion? 2) do newly inserted TEs adopt the methylation pattern of their genomic context? and 3) do new TE insertions affect the methylation of their flanking regions? While most non-TE insertions exhibit DNA methylation patterns consistent with their genomic context, TE insertions are generally highly methylated, exhibiting distinct, class-specific patterns, and with profound variation within TE bodies. A small percentage of *Alu* insertions are hypomethylated, particularly those inserted within hypomethylated CpG islands. By comparing DNA methylation of flanking regions of TE insertions between individuals with and without the TE insertions, we revealed that majority of TEs exhibited minimal impact on nearby regions, although numerous exceptions exist where the methylation status of both L1 and *Alu* insertions “leak” into nearby regions, leading to either methylation spreading or hypomethylation sloping shores. In conclusion, we demonstrated the methylation calling capability of third-generation sequencing and its unique advantage in characterizing epigenomic features within non-reference positions. While TE insertions primarily exhibit methylation patterns restricted within their boundaries, some TEs are able to engage in context-dependent complex interactions with genomic neighborhood.

## INTRODUCTION

Cytosine methylation at CpG sites is prevalent in vertebrate genomes. The majority of CpG sites in the human genome are methylated in most somatic tissues (Bird 1986; The ENCODE Project Consortium 2012; Roadmap Epigenomics Consortium et al. 2015). It is generally believed that the CpG methylation is a repressive mark commonly associated with heterochromatin and transposable elements (TEs) silencing, but is absent from promoter of active genes (Moore et al. 2013). In addition, methylation pattern is dynamic during development and disease progression (Jones 2012; Smith et al. 2024). During evolution, methylation-mediated deamination converts methylated cytosine to thymine, causing the general depletion of CpG sites in most vertebrate genomes (Bird 2002). However, there are CpG islands (CGIs) in the genome that are relatively enriched with GC content and CG motif (generally defined by sequences > 200bp with >=50% GC content and the observed/expected CpG ratio >=0.6) (Fazzari and Greally 2004). It is believed that they represent conserved unmethylated regions in germ cells during development (Jones 2012; Smallwood et al. 2011) and many were found in the promoters of conserved genes (Sved and Bird 1990; Han et al. 2008).

Given the importance of CpG methylation in genome regulation and evolution, considerable efforts have been devoted to characterizing cytosine methylation. Microarray based methylation chip, MeDIP-seq, MRE-seq, and RRBS-seq are all popular options to interrogate methylation states of different CpGs across the genome (Hodges et al. 2009; Weber et al. 2005; Meissner et al. 2008; Maunakea et al. 2010). Because of its universal coverage across the genome, Whole Genome Bisulfite Sequencing (WGBS) has been considered the gold standard in cytosine methylation calling since its debut in the 2nd generation sequencing (shotgun based sequencing of several hundred base pairs) era (Lister et al. 2009; Cokus et al. 2008; Laurent et al. 2010).

Despite its many advantages, WGBS has several limitations. For example, the bisulfite treatment that chemically distinguishes methylated from unmethylated cytosine damages the DNA molecules and complicates the library preparation before sequencing. In addition, WGBS typically relies on identifying mismatches between bisulfite-treated reads and the reference genome to call methylation, making methylation calling of cytosines of non-reference regions non-trivial. There are new methods developed recently to improve WGBS but ultimately, they are limited by the 2nd generation sequencing (Vaisvila et al. 2021; Nunn et al. 2022).

The 3rd generation sequencing technologies, pioneered by PacBio and ONT, have greatly matured in recent years (van Dijk et al. 2018; Sigurpalsdottir et al. 2024). This improvement results in much longer reads and the ability to distinguish modified bases, included methylated cytosine, during basecalling (Flusberg et al. 2010; Schatz 2017; Stergachis et al. 2020; Xu and Seki 2020). Thus, methylation calling and sequencing can be done simultaneously, making cytosine methylation calling much simpler and more accessible without resort to specific chemical treatment to target cytosine methylation. Despite the promising outlook, the signal changes caused by methylation is usually small and variable, and it is not until the recent development of deep learning models that CpG methylation calling on SMRT and Nanopore data has seen dramatic improvement in accuracy (Simpson et al. 2017; Tse et al. 2021; Ni et al. 2023; Liu et al. 2021).

More than just providing better mapping and coverage to highly repetitive regions, the 3rd generation sequencing enabled us to identify non-reference insertions along with their methylation status at individual level including generally methylated TEs. However, there are many outstanding questions regarding the epigenetic response to newly inserted TEs. Which of them are methylated, which of them are not? Is their methylation being influencing by genomic regions into which they insert? Can they affect the methylation status of nearby genomic regions? If they can, how far away would their influence continue? The advance in 3rd generation sequencing makes it much easier to directly address these questions at scale. Recently, third generation sequencing was used to characterize the methylation profile of repetitive elements with high resolution (Ewing et al. 2020; Gerdes et al. 2023; Bodea et al. 2024).

It has been reported in mice that a newly inserted hypermethylated TE can spread methylation from the insertion site to the flanking regions until blocked by other regulators (Turker 1999, 2002), resembling the spreading of heterochromatin (Sentmanat and Elgin 2012). The spreading of TE methylation can further influence the host genome during evolution by altering mutation pattern (Zhou et al. 2020). Not only hypermethylation can spread out of TEs, tissue-specific hypomethylated CGIs introduced by new L1 insertions can also create so-called “sloping shores” in otherwise hypermethylated regions (Grandi et al. 2015). There were also studies reporting the methylation of new TE insertions have limited effect on nearby gene expression pattern in plants (Choi and Purugganan 2018; Wyler et al. 2020). Recently, Lanciano et al. investigated the local impact of L1 insertion using bsATLAS-seq and ONT reads in multiple cell lines and found regions 300bp upstream of L1 insertions can be influenced by the methylation status of L1 (Lanciano et al. 2024).

Taking advantage of the high coverage of the Human Pangenome Reference Consortium (HPRC) 3rd generation sequencing data and latest methylation calling methods, we compared the whole-genome CpG methylation in five human lymphoblastoid cell lines (LCLs) using Single Molecule, Real-Time (PacBio), Nanopore (ONT), and WGBS technologies. After benchmarking the accuracies of methylation calling of the 3rd generation sequencing, we investigated DNA methylation profiles of polymorphic TE insertions, the relationship between DNA methylation patterns of the insertions and their flanking genomic regions, as well as how local methylation could be affected by the new insertions. We found that the 3rd generation sequencing is capable of accurately characterizing CpG methylation at non-reference positions that are inaccessible with conventional WGBS and determined DNA methylation of polymorphic TE insertions and how they interact with their flanking regions.

## RESULTS

### Both PacBio HiFi and Nanopore ONT reads provide accurate DNA methylation estimation

The Human Pangenome Reference Consortium (HPRC) published the first human pangenome reference graph representing 47 genetically diverse individuals (Liao et al. 2023). Out of 47, the DNA methylation status of 32 were determined using PacBio HiFi reads (download). In addition to the rich genetic information and various annotations, the HPRC resources also enabled direct methylation analysis using long read sequencing data. Taking advantage of these resources, we performed additional CpG methylation calling using ONT reads (R9.4.1) from five individuals who were part of the first draft of the Human Pangenome Reference (HG00741, HG00621, HG01952, HG01978, and HG03516) and Whole Genome Bisulfite Sequencing (WGBS) of these same five individuals (Methods). After calling CpG methylation, we aligned all reads to the human reference genome hg38 (Fig. 1a).

**Figure 1.**
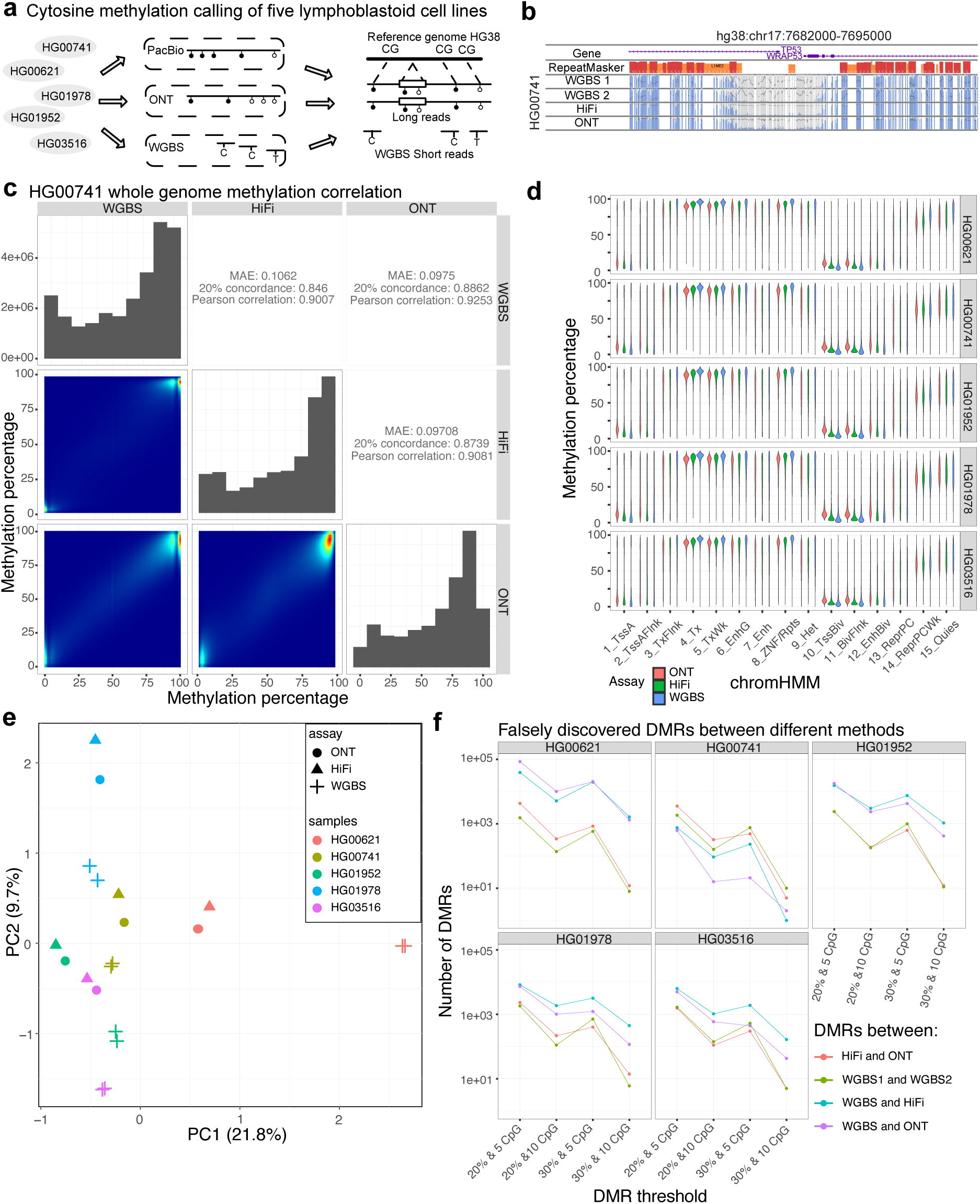
**a**: Study design of methylation calling benchmarking. We performed CpG methylation calling for five HPRC samples (HG00621, HG00741, HG01952, HG01978, and HG03516), using PacBio, ONT, and WGBS. We mapped all reads to the human reference genome hg38 to compare their differences. **b**: WashU Epigenome Browser screenshot of the two WGBS replicates, HiFi, and ONT methylation of HG00741 on the 10 kb window around the TP53 gene promoter. All methylC tracks Y-axis percentage was from 0% to 100%. **c**: Whole genome methylation percentage correlation between WGBS, HiFi, and ONT methylation of HG00741. The whole genome methylation percentage distribution was represented as a density plot. The correlation heatmap and metrics (Mean Average Error, 20% concordance, and Pearson correlation) were plotted on the left-bottom and right-top, respectively. **d**: Methylation percentage distribution of the GM12878 15 chromHMM states across all five samples. The 15 chromHMM states were listed on the X-axis and methylation calling results from different methods were colored differently. **e**: PCA plot of the top two principal components of the whole genome methylation from the five HPRC samples. For each sample, we performed two replicates of WGBS in addition to HiFi and ONT methylation calling. We used different shapes to represent different methods (circle, triangle, and cross for ONT, HiFi, and WGBS) and different colors to represent different samples (red, yellow, green, blue, and purple for HG00621, HG00741, HG01952, HG01978, and HG03516, respectively). **f**: Number of falsely discovered differentially methylated regions (DMRs) using different threshold cut-offs. Four different thresholds (20% average methylation difference with at least 5 CpG sites, 20% average methylation difference with at least 10 CpG sites, 30% average methylation difference with at least 5 CpG sites, and 30% average methylation difference with at least 10 CpG sites) were represented on the X-axis. The number of DMRs was represented on the Y-axis. The number of DMRs between different methods under different conditions was plotted using different colors.

We first compared DNA methylation calls between HiFi, ONT, and WGBS for all CpG sites in the reference genome hg38. For HiFi reads, we adopted the deep learning based mathematical model (PB-model) developed and suggested by PacBio to estimate methylation percentage of individual CpG sites (https://github.com/PacificBiosciences/pb-CpG-tools). We calculated mean average error (MAE), 20% concordance, and Pearson correlation between WGBS, HiFi, and ONT for each sample in pairwise comparisons (HG00741 comparison in Fig. 1c and the other four in Supplemental Fig. S1a-d). We found both HiFi and ONT methylation calls highly correlated with their matching WGBS results (MAE: 0.1062 and 0.0975; 20% concordance: 0.846 and 0.886; Pearson correlation: 0.901 and 0.925 for HiFi and ONT, respectively for HG00741) in all five samples (Fig. 1c and Supplemental Fig. S1a-d). We examined the distribution of methylation level across different genomic contexts (defined by the GM12878 chromHMM states) and found strong concordance among WGBS, HiFi and ONT across all five samples (Ernst and Kellis 2012) (Fig. 1d). We then used the WashU Epigenome Browser to illustrate the comparisons between WGBS, HiFi and ONT (Fig. 1b).

To gain a global view of the difference and similarity across methods, we performed Principal Component Analysis (PCA) and plotted the first two Principal components in a 2D plot for visualization (Fig. 1e). In all five samples, the two matching WGBS replicates consistently clustered together; HiFi and ONT methylation results from the same samples also clustered together. The PCA distance between WGBS and the long read methylation results seemed to be larger than the distances between HiFi and ONT.

To better understand the scale of differences between the different methods, we defined both differentially methylated loci (DMLs) and differentially methylated regions (DMRs) between HiFi, ONT, and WGBS, as well as between two WGBS replicates of the same sample (Wu et al. 2015). Since the DMRs here were defined using the same samples, they represented the false positive discoveries, and allowed us to estimate an empirical false discovery rate. Using different thresholds, we found that the numbers of falsely discovered DMRs between HiFi and ONT were comparable with the numbers derived from the two WGBS replicates (Fig. 1f). Thus, the methylation calling differences between HiFi and ONT was quite small, generally comparable with the differences we observed between the two WGBS replicates.

To further understand the methylation differences between long-read and short-read methods, we examined DMLs and DMRs called between the two methods. We found that the DMLs between long and short reads that exhibiting a >50% methylation difference were predominantly contributed by CpG sites with abnormally high short-read coverage (>50X) (Supplemental Fig. S1e), suggesting short-read mapping in these regions were more challenging than others. In contrast, long-read sequencing coverage or GC content did not correlate with falsely discovered DNA methylation differences (Supplemental Fig. S1f,g). We also found that low complexity and simple repeats contributed to a higher fraction of DMRs with a methylation difference >40% (Supplemental Fig. S1h). Since these regions also suffer more from imprecise mapping of short reads, we reasoned that the small discrepancy between long-read and short-read based methods in assessing DNA methylation was systematic and likely a result of reference mapping bias (Pollard et al. 2018). A systematic benchmarking analysis with a focus on alignment bias will be required in the future to fully understand this phenomenon.

### The methylation patterns of TE insertions

Because the 3^rd^ generation methylation calling was conducted on raw reads without aligning to the reference genome, it enabled methylation calling of non-reference regions previously inaccessible with 2nd generation sequencing based methods (at least not trivially). Thus, we investigated the methylation status of non-reference insertions with a focus on polymorphic TE insertions and their correlation with the methylation levels of their flanking regions. Here we took advantage of the recent release of the draft Human Pangenome Reference made available by the HPRC, which included HiFi reads with methylation calling from 32 HPRC Year1 samples (download) as well as their genetic variations (download) (Liao et al. 2023). We further annotated insertions combining the Transposable Element Insertion Annotation pipeline MELT-LRA, PALMER2 and xTEA (Methods) (Gardner et al. 2017; Zhou et al. 2024; Chu et al. 2021). We classified TE insertions into four categories: *Alu*, L1, SVA, and HERVK. In total, we identified 6,752 *Alu*, 410 L1, 334 SVA, 16 solo LTR5 and 9 HERVK insertions from the 32 HPRC samples (Supplemental Table S1). For convenience, we termed these TE insertions non-reference TEs, and TEs annotated in hg38 reference TEs.

Using custom pipelines (Methods), we extracted non-reference TEs and the HiFi reads derived from these regions (Fig. 2a). We first assessed if we had sufficient long-read coverage on non-reference TEs. We calculated the relative CpG site coverage of each TE insertion compared with the genome-wide average coverage based on HiFi data and observed the two expected peaks for each TE class, representing coverages of heterozygous and homozygous insertions (Supplemental Fig. S2a). The average genome-wide coverage was 45.9X, thus we believe that the coverage on TE insertions was sufficient for the purpose of our analysis. We also extracted HiFi methylation of reference TEs for comparison (Fig. 2a).

**Figure 2.**
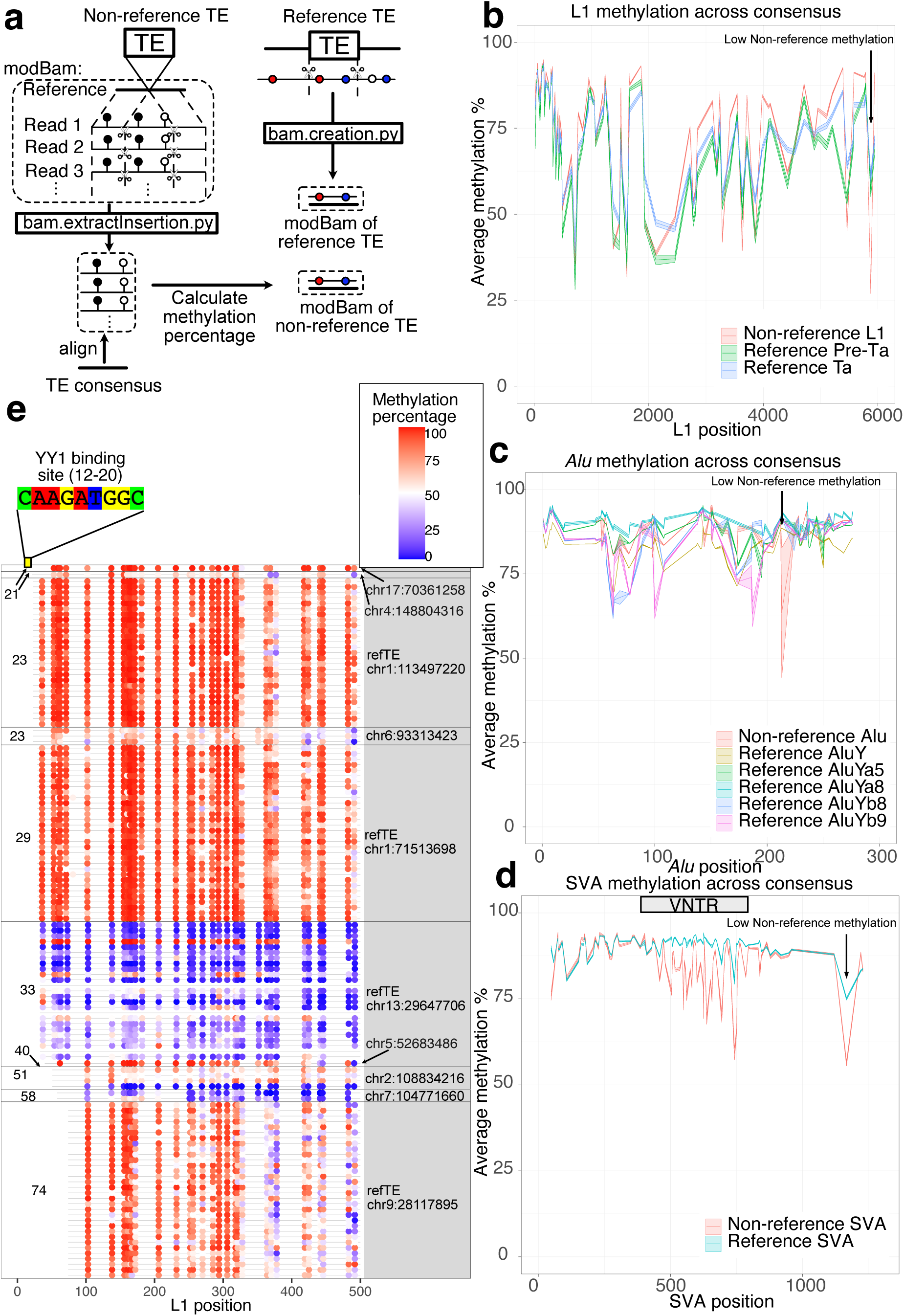
**a**: Schematic representation of our pipeline to extract reference and non-reference insertions along with their methylation annotation. We developed a script (bam.extractInsertion.py) to extract truncated DNA sequences from reads while retaining positional methylation tags (subseq modBam file). We then aligned this modBam to the corresponding TE consensus and calculated the methylation profile of the TE insertion. We also extracted sequences from the reference genome and the methylation percentage from the bedMethyl file to create another modBam file of reference TEs and their methylation (bam.creation.py). The methylation would be anchored to the TE consensus after alignment. **b**: Methylation profile of L1 insertions. X-axis represented positions on the L1Hs consensus, Y-axis represented methylation percentage with 95% standard error of the mean illustrated as shaded areas. Reference L1pre-Ta, reference L1Ta, and non-reference L1 insertions were labeled with different colors. Only full-length elements (truncation < 50bp on both ends) were included for profiling. **c**: Methylation profile of *Alu* insertions with 95% standard error of mean. X-axis represented positions on the *AluY* consensus, Y-axis represented methylation percentage. *AluY*, *AluYa5*, *AluYa8*, *AluYb8*, *AluYb9* on the reference and non-reference *Alu* insertions were labeled with different colors. **d**: Methylation profile of SVA insertions with 95% standard error of mean. X-axis represented positions on the SVA_F consensus, Y-axis represented methylation percentage. Reference SVA and non-reference SVA insertions were labeled with different colors. The VNTR of SVA consensus were labeled on the graph with a gray box. **e**: The first 500bp methylation of L1 insertions whose first 20-50bp were truncated. Start position was labeled on the left, and the insertion coordinates on hg38 were labeled on the right. The “refTE” indicates it was an L1 insertion on the hg38 reference. Each row represented one of the 32 samples. The methylation percentage of each CpG site was illustrated by the blue-red (low to high) color scale.

For the five HPRC individuals with WGBS, HiFi and ONT methylation results, we phased both HiFi and ONT reads to maternal and paternal assemblies and mapped them onto maternal or paternal haplotypes, respectively (download) (Patterson et al. 2015; Holt et al. 2024). Using the reference-assembly alignment, we created comparative epigenome browser datahubs to interactively examine the methylation of non-reference insertions (HiFi datahub: https://epigenomegateway.wustl.edu/browser2022/?genome=hg38&hub=https://wangcluster.wustl.edu/∼xzhuo/hifi_methylation/hifi.all.json; ONT datahub: https://epigenomegateway.wustl.edu/browser2022/?genome=hg38&hub=https://wangcluster.wustl.edu/∼xzhuo/hifi_methylation/ont.all.json).

We first compared DNA methylation profiles of full-length non-reference TE insertions with their counterparts in the reference genome. We extracted *52,802 Alu*Y, 2,746 *Alu*Ya5, 36 *Alu*Ya8, 2,111 *Alu*Yb8, and 301 *Alu*Yb9 from hg38 using repeatMasker annotation. Similarly, we extracted 124 SVA_E, 331 SVA_F, and 481 LTR5 as reference TEs. We also downloaded 146 active L1 on hg38 from L1 Base 2 (Penzkofer et al. 2017). Active L1s were further classified into L1Ta and L1Pre-Ta using the defining variant at position 5932 (A as L1Ta and G as L1pre-Ta) based on L1base 2 (Boissinot et al. 2000). By anchoring methylation to a consensus-based alignment, we significantly reduced the noise in methylation signals compared to previous studies (Ewing et al. 2020; Gerdes et al. 2023). This allowed us to observe sharp methylation fluctuations between adjacent CpG sites within transposable elements (TEs) (Fig. 2b-d). Our findings suggested that certain CpG sites, such as position 5875 in L1HS, were consistently lowly methylated. Overall, DNA methylation profile of reference and non-reference TE insertions showed high methylation levels and were similar in all four classes (*Alu*, L1, and SVA profile in Fig. 2b-d and LTR5 profile in Supplemental Fig. S2b). Interestingly, a single CpG site in each of *Alu*, L1 and SVA (at position 213, 5875, 1166 respectively) exhibited statistically significant differences between non-reference insertions and reference copies. Also profoundly, the internal CG rich, variable number of tandem repeats (VNTR) region of SVA was methylated lower in the non-reference insertions compared with their counterpart in the reference copies (Fig. 2d). The overall methylation plots for all but *Alu* insertions (which were too numerous to display individually) were available in Supplemental Fig. S2c-e.

To our knowledge, the relatively low methylation of VNTR within SVA elements have never been reported before. Ewing et al. found that the VNTRs are methylated higher than its flanking regions in SVA (Ewing et al. 2020) and we confirmed their observation within reference SVAs (Supplemental Fig. S2d). However, the methylation of VNTR of non-reference SVAs in our LCL samples were noticeably lower than both their flanking regions and their counterparts in the reference genome (Fig. 2f), and it resembled the methylation of mouse L1 5’UTRs with different length of monomers (Gerdes et al. 2023). Aligning full-length SVAs, often >2kb long, to the 1.3kb consensus may create some alignment issue, especially in the VNTR region. However, we do not believe this could explain our observation: the long read methylation calling was done during basecalling process, independent of alignment, and the presence of many lowly methylated CpGs was visible across many non-reference SVAs (Supplemental Fig. S2f).

The transcription factor YY1 is an important regulator of L1 activity in mammals (Saha et al. 2024). It was reported that L1 insertions missing the YY1 binding site, located at positions from 12 to 20 in the L1HS, could escape 5’ methylation in a tissue-specific manner (Sanchez-Luque et al. 2019). It was also proposed that YY1 binding affects the methylation of L1 in a cell type-specific manner (Lanciano et al. 2024). To investigate whether the YY1 binding site could affect polymorphic L1 methylation in LCL cells, we examined the 5’ end methylation of 10 L1 insertions whose first 20-100bp were truncated (Fig. 2e).

The YY1 motif is located at position 12-20 in L1 consensus and none of the 10 L1s here contained the YY1 binding site. Despite that, we found five loci, missing 20 bp to 28 bp from the 5’end, were hypermethylated at the L1 promoter. In contrast, three out of the five loci with >32 bp truncated at the 5’ end were hypomethylated. Two loci, missing 39 and 73 bp respectively, remained hypermethylated. Our data thus suggest that missing the YY1 binding site is not sufficient for the demethylation of L1 promoter, at least not in LCL. Interestingly, we found individual DNA methylation differences of the same L1 insertions. The reference L1 insertion with the first 31 bp at the 5’end truncated at chr13:29641706-29647706, denoted as Chr13delta31 in Sanchez-Luque et al. (Sanchez-Luque et al. 2019), displayed variation among our 32 samples: its promoter was hypomethylated in most of the samples but hypermethylated in HG02717 (Supplemental Fig. S2g).

### Low methylation correlation between TE insertions and their flanking regions

We next investigated the relationship between DNA methylation of the insertions and their flanking regions. We compared the average methylation levels of non-reference insertions, categorized based on their TE annotation, with the methylation levels of their 300bp flanking regions on both 5’ end and 3’ end (Fig. 3a). TE insertions, particularly *Alu* and SVA insertions, exhibited high methylation levels regardless of the methylation levels of their flanking regions (Fig. 3b). On average, we observed weak correlations between TEs and their flanking regions (linear regression slope < 0.2, R^2^ ∼ 0.1). We did not calculate the correlation of HERVK insertions because of the insufficient number of non-reference HERVK insertions. This pattern was in sharp contrast to non-TE insertions, whose methylation levels were highly correlated with their nearby flanking regions (slope = 0.88, R^2^ = 0.61) (Fig. 3b). The pattern for non-TE insertions did not change much when we separated them based on whether they contained tandem repeats, whether they were in CpG islands, or their CpG density (Supplemental Fig. S3a). Reference TE insertions displayed a similar pattern to that of non-reference TEs (Supplemental Fig. S3b). Among reference TE insertions, LTR5 exhibited the highest linear correlation slope of approximately 0.5 (Supplemental Fig. S3b).

**Figure 3.**
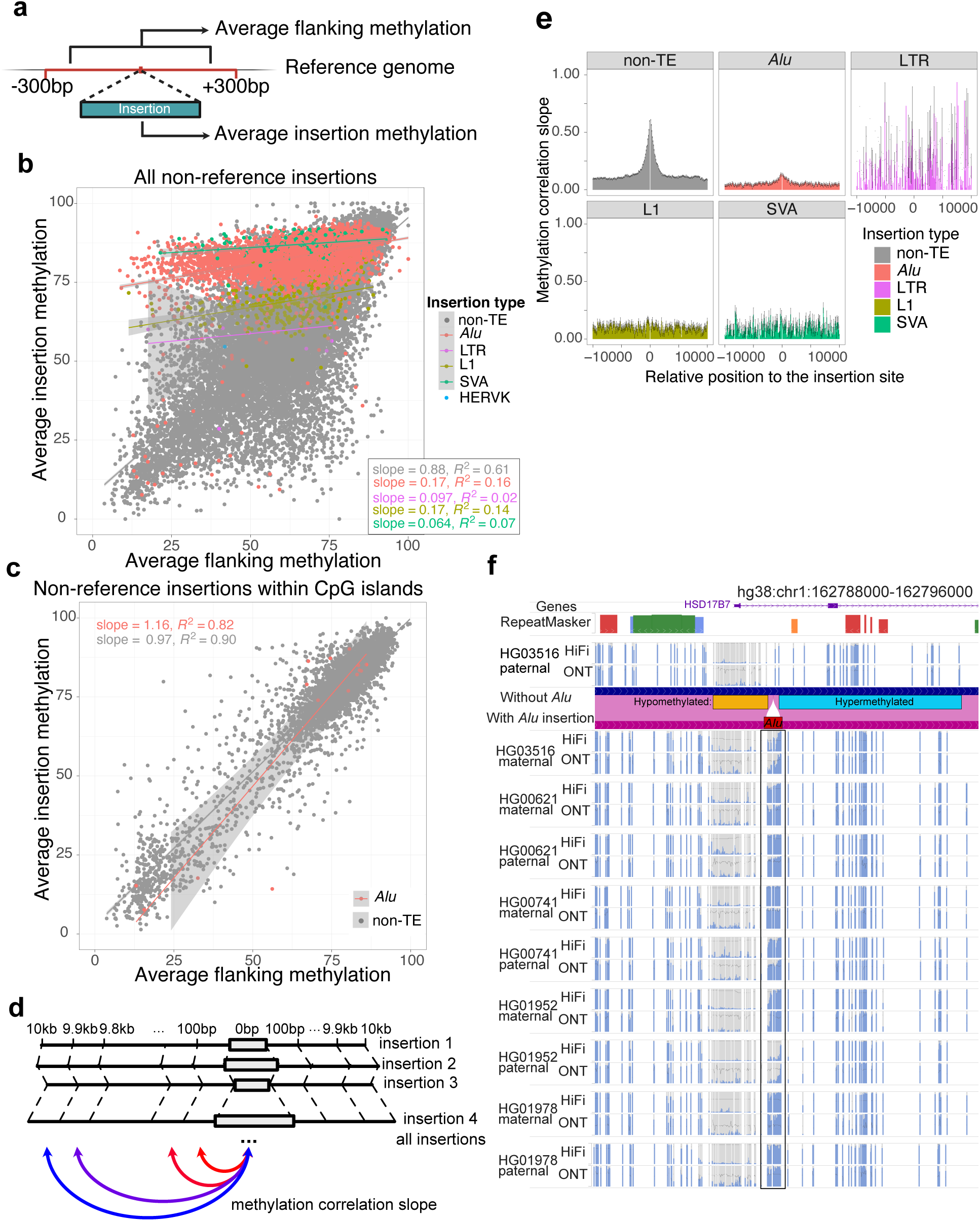
**a**: Schematic representation of average methylation calculation of all insertions and associated 300bp flanking regions. **b**: Methylation correlation between insertions and 300bp flanking regions. The average methylation of insertions was represented on the Y-axis, while the methylation of flanking regions on the X-axis. *Alu*, LTR, L1, SVA, and non-TE insertions were represented by red, green, blue, and purple dots, respectively, whereas non-TE insertions were labeled grey, serving as a background. Linear regression lines with standard errors were also plotted and labeled with same color scheme. **c**: Methylation correlation between insertions within CpG islands and 300bp flanking regions. The average methylation of insertions was represented on the Y-axis, while the methylation of flanking regions on the X-axis. *Alu*, and non-TE insertions were represented in the same way as they are in 3b. **d**: Schematic representation of insertion-flanking methylation regression as a function of insertion-flanking distance. For each distance from 100bp to 10kb, we performed linear regression between insertion and flanking average methylation and estimated the linear regression slope and standard errors. **e**: Linear regression slope between insertion and flanking methylation as a function of the insertion-flanking distance from 10kb upstream to 10kb downstream. Standard errors were represented by error bars. *Alu*, LTR, L1, SVA and non-TE insertions were plotted in separated plots with different colors. **f**: The WashU Epigenome browser views of an *Alu* insertion between hypomethylated and hypermethylated regions. The insertion was absent from HG03516 paternal haplotype but present in all the other nine haplotypes. A genome-align track was used to illustrate the insertion site. The *Alu* insertion methylation was boxed, showing a gradual change of methylation from left to right in some of them. Both HiFi and ONT methylC tracks displayed methylation percentage from 0%-100%.

CpG island (CGI) is an important genomic feature, and its methylation status plays a significant role in gene regulation^1^. When we restricted the above analysis to non-TE insertions within CGIs, we observed a stronger correlation between the insertions and their flanking methylation (slope = 0.97, R^2^ = 0.90) (Fig. 3c). Remarkably, this correlation extended even to *Alu* insertions within CGIs. In contrast to the general hypermethylation of *Alu* insertions genome-wide, the methylation of *Alu* insertions within CGIs correlated highly with their flanking methylation (slope = 1.16, R^2^ = 0.82) (Fig. 3c).

Despite the majority of *Alus* being hypermethylated, a small percentage were hypomethylated. For instance, at chr11:59565869, a polymorphic *Alu* insertion within a hypomethylated CpG island was hypomethylated. We plotted the methylation of this *Alu* along with its 300bp flanking regions across all 32 samples (Supplemental Fig. S3c). The insertion site was hypomethylated in all samples, and the *Alu* insertion itself appeared to have adopted the hypomethylation status in all five samples in which it was present. This region can be further examined in sample HG03516, whose *Alu* insertion was heterozygous (Supplemental Fig. S3d). The regional hypomethylation, presence of the *Alu* insertion, were supported by HiFi, ONT, and WGBS analysis (Supplemental Fig. S3d,e).

Several *Alu* elements exhibited a methylation slope that connected hypomethylated and hypermethylated regions. We illustrated four such cases in Supplemental Fig. S3f. At chr1:162791517, chr2:218333959, and chr9:71308789, the 300bp upstream of the insertion was hypermethylated, while the downstream region was hypomethylated. In contrast, at chr4:163587802, the upstream region was hypomethylated, while the downstream region was hypermethylated. In each of these four cases, the *Alu* insertion displayed a gradual transition between hypo- and hypermethylation, with some variation observed among samples. We displayed the antisense *Alu* insertion at chr1:162791517 in our comparative browser by phasing reads to paternal and maternal alleles (Fig. 3f). The ONT and HiFi data showed high agreement with each other on methylation level of this non-reference insertion. Downstream of the *Alu* was hypomethylated (left side), while upstream of it (right side) was hypermethylated in all samples. Interestingly, there were variations among individuals and between haplotypes at this locus. While most instances of this insertion were highly methylated throughout the *Alu* element, the methylation level in the HG03516 maternal allele exhibited a transition from low to high in the middle of the element. In contrast, this insertion was hypomethylated in the HG01952 paternal allele. No other sequence feature seemed to be associated with the methylation pattern difference across individual alleles around chr1:162791517.

Our analysis thus far has revealed a strong correlation in DNA methylation levels between non-TE insertions and their 300bp flanking regions, and a much weaker correlation when the insertions were TEs. We next investigated the insertion-flanking methylation correlation beyond 300 bp as a function of the distance between the insertion and the nearby regions. For each non-reference insertion, we calculated DNA methylation levels in its 10 kb flanking regions on both directions at a resolution of 100-bp non-overlapping windows (Fig. 3d). We grouped the average methylation of all 100-bp windows from different insertions by their distance from the insertion sites and performed linear regression between insertion and flanking regions. Since the slopes of the linear regression can reflect the methylation correlation between these regions, we displayed the slope of the linear regression as a function of the distance between the flanking windows and the insertion (Fig. 3e). This analysis revealed that the methylation correlation slope between non-TE insertions and their flanking windows quickly faded away as the distance between the flanking window and insertion site increased, with an estimated elbow point being ∼2.5kb away from the insertion. The slope decayed to and stayed at around 0.1 when the distance reached 5kb and beyond. TEs once again displayed a strikingly distinct pattern. There was no observable correlation between L1, SVA, and HERV insertions and their flanking regions. We did however observe a weak correlation between *Alu* insertions and their flanking regions but it also quickly faded away (Fig. 3e). This observation could be confounded if TE insertions were longer than non-TE insertions. To rule out this possibility, we investigated how the size of non-TE insertion could affect the insertion-flanking methylation correlation. If the methylation correlation is merely a function of the insertion size, we would expect ∼300bp and ∼1000bp non-TE insertions behave similarly to *Alu* and LTR insertions, for example. We categorized non-TE insertions by their sizes into five bins, then calculated the regression slope in each bin (Supplemental Fig. S3g). Despite a slightly decrease of the peak methylation correlation as a function of increased non-TE insertion size, the correlation was much higher than that of TE insertions of comparable length (for example, the slope distribution of 300-1000bp non-TE insertions was much higher than that of *Alu* and LTR insertions). This result suggests that the methylation of non-TE insertions, but not TE insertions, are highly correlated with that of their flanking regions.

### Most TE insertions do not change the methylation level of their flanking regions

Some non-reference insertions could be deletion events in the human population (Lee et al. 2020b). However, TE insertions are essentially homoplasy-free (Batzer and Deininger 2002) and represent recent polymorphic insertions in the human population. Therefore, they are excellent candidates to investigate how new insertions might have influenced the methylation of their inserted regions.

Previous studies have suggested that repressive chromatin marks including DNA methylation deposited on newly inserted TEs can spread beyond TE/genome boundary (Turker 2002; Sentmanat and Elgin 2012; Zhang et al. 2012). It was also reported that hypomethylated L1 could drag down the methylation at its 5’ end regions and create “sloping shores” (Grandi et al. 2015). Recently, Lanciano et al. found L1 insertions can have an impact on the methylation of CpG sites within 300bp upstream based on both bs-ATLAS-seq and ONT sequencing of multiple cancer cell lines (Lanciano et al. 2024).

The Human Pangenome Reference Consortium (HPRC) resources provided a unique opportunity to investigate the impact of TE insertions on nearby regions. For each polymorphic non-reference insertion relative to hg38, we separated HiFi reads from the 32 HPRC Year1 samples into two groups: alleles with the insertion and alleles without the insertion (Fig. 4a). We then calculated the average methylation level of the inserted regions (insertion), the 300bp flanking regions of the insertion from both ends (insertion flanking), and the same flanking regions of the allele without insertion (empty flanking) (Fig. 4a).

**Figure 4.**
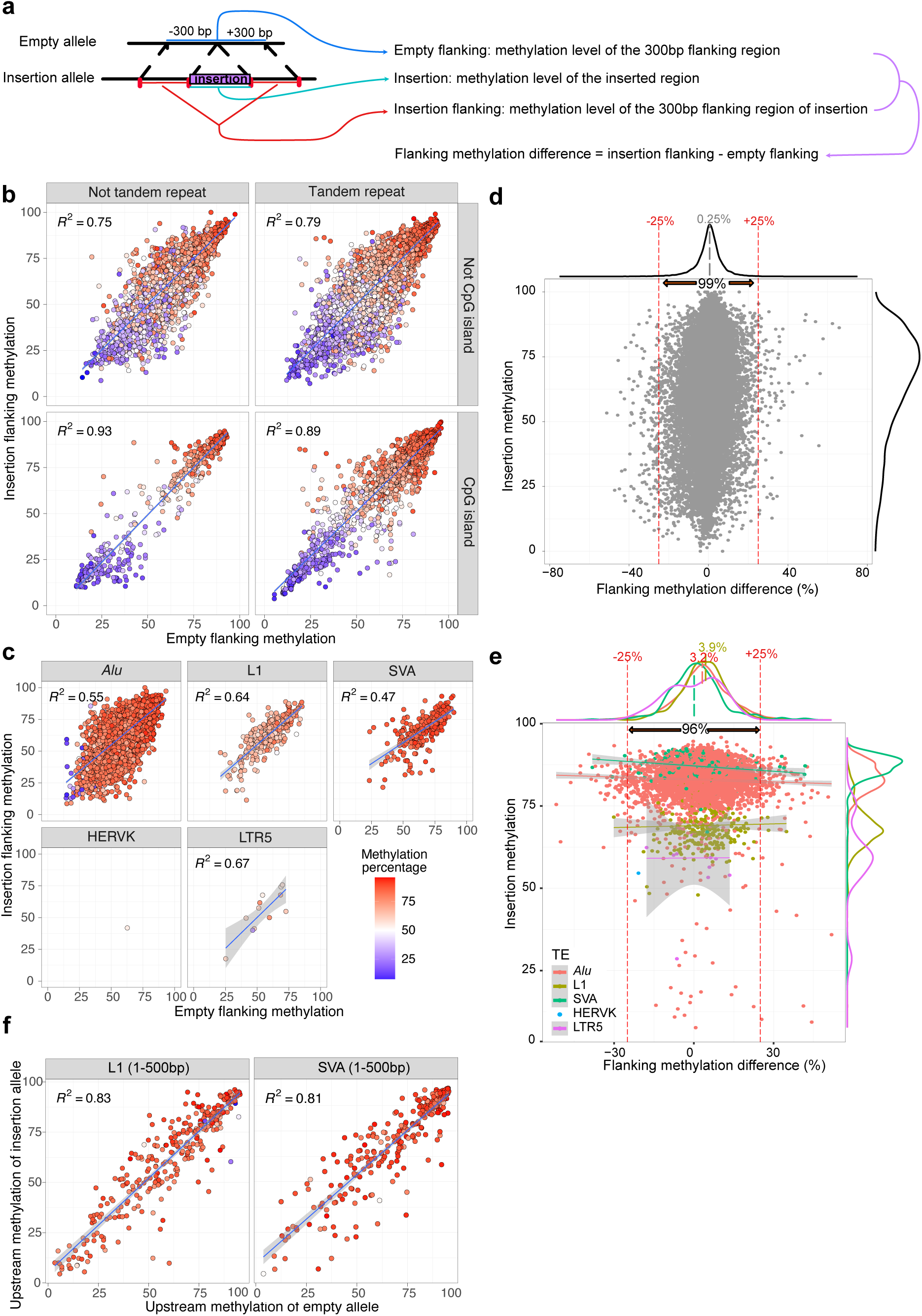
**a**: Schematic representation of a polymorphic insertion with aligned long reads. The allele without the insertion site (top) and the allele with the insertion site (bottom) were represented as reference allele and insertion allele. The average methylation level of the insertion, insertion flanking regions, and empty flanking regions were calculated from reads from all 32 samples. We then calculated flanking methylation difference by subtracting the methylation level of empty flanking region from that of insertion flanking region. **b**: Methylation level of empty flanking (X-axis), insertion flanking (Y-axis), and non-TE inserted regions (blue-red color scale as shown in Fig. 4c). Insertions were separated into four facets by whether they are tandem repeats and whether they are CpG islands. Linear regression R^2^ values were shown on each panel. **c**: Similar to Fig. 4b, but the methylation level of different TE insertions was presented and separated by the TE insertion types. R^2^ were displayed in the top left corner on each panel. **d**: Dot plot illustrated the relationship between methylation level of insertion (Y-axis) and methylation difference between empty and insertion alleles of the flanking regions (X-axis). The average methylation difference of all non-TE insertions was labeled at 0.25%. Red dotted lines indicated the methylation difference threshold of +/-25%, and 99% of non-TE insertions fall within this range. The insertion methylation density was displayed on the right side of the figure. **e**: TE insertion flanking methylation differences plotted in the sample way as in Fig. 4d. Different TE insertions were distinguished by different colors. The percentage of all insertions within 25% differences and the average flanking methylation differences of different TEs were labeled. The insertion methylation density was shown on the right of the figure. The average methylation of each TE class was labeled with the same color scheme. **f**: Methylation level of 300 bp upstream of the empty allele (X-axis), 300 bp upstream of the insertion allele(Y-axis), and the first 500 bp inserted methylation (blue-red color scale) of L1 and SVA insertions displayed similar to Figure 4b and c.

To illustrate the relationship between these three regions, we plotted the methylation levels of insertion flanking and empty flanking on the X-Y axis using a dot plot. We colored the dots based on the methylation level of insertions. We separated non-TE insertions by whether they were annotated as tandem repeat or CGIs, and classified TE insertions by TE classes. In general, the methylation levels of insertion, insertion flanking, and empty flanking correlated with each other for non-TE insertions (Fig. 4b). However, for TE insertions, the flanking methylation levels of insertion and empty sites showed a strong correlation independent of the methylation of the insertion (Fig. 4c).

To better understand the subtle changes in flanking methylation levels between insertion and empty sites, we defined the flanking methylation difference for each insertion by subtracting the flanking methylation of the empty site from the flanking methylation of the insertion site (Fig. 4a). We reasoned that an allele without a TE insertion represents the ancestral allele^56^. If the insertion increases flanking methylation, the difference would be positive; if it decreases methylation, the difference would be negative. Conversely, if the insertion has no impact on flanking region methylation, the difference would be zero.

We first plotted the correlation between flanking methylation variations and the methylation of non-TE insertions (Fig. 4d). The overall methylation differences between orthologous flanking regions with or without insertions were predominantly centered around zero (population average was 0.25%) – 99% of the data points (28344/28621) fell within the −25% to +25% range. Notably, these variations were independent of the methylation level of the insertion itself. However, there indeed existed multiple instances where hypermethylated insertions were associated with increased flanking methylation, and hypomethylated insertions were linked to decreased methylation (Fig. 4d). Collectively, these findings strongly indicate that, although there are noted exceptions, non-TE insertions generally do not significantly impact flanking region methylation.

We then investigated the same relationship in the context of TE insertions (Fig. 4e). Compared to non-TE insertions, the range of methylation differences was slightly wider (96%, 5075/5263, within −25% and 25%), indicating significantly increased variations in flanking region methylation associated with TE insertions (F-Test to Compare Two Variances: p-value < 2.2e-16). On average, flanking methylation differences centered around zero for LTR and SVA insertions. However, the average flanking methylation differences for *Alu* and L1 insertions were significantly larger than zero (3.2% and 3.9%, respectively). Furthermore, there were more *Alu* and L1 insertions with methylation increase more than 25% than decrease more than 25% (112 vs 62, and 4 vs 1, respectively). The range of methylation differences we observed was quite large. For example, the mean methylation level of inserted *Alus* was 82.7%, and *Alu* insertions with methylation level greater than this mean value were associated with a flanking region methylation difference ranging from −42.74% to +42.66%, with 21 of them associated with an increased flanking region methylation by greater than 30%. Similarly, the mean methylation level of inserted L1s was 69.2%, and L1 insertions with methylation level greater than this mean value were associated with a flanking region methylation difference ranging from −30.1% to +27.8%, but none of them was associated with an increased flanking region methylation by greater than 30%. These examples only weakly support the hypothesis that DNA methylation deposited on newly inserted TEs could spread and increase local DNA methylation level. Contrary to this conventional view, our genome-wide assessment suggests that the spreading is limited, and those more drastic influence of TEs on nearby genomic methylation may be an exception rather than the rule.

It has been reported that L1 promoter methylation can influence the nearby upstream methylation (Lanciano et al. 2024). To test this hypothesis using our data, we created a dotplot similar to Fig. 4c, but using the 5’end 500bp of L1/SVA insertions along with the 300bp upstream region genomic sequences (Fig. 4f). Notably, despite the existence of a few exceptions, the upstream methylation of empty sites consistently matched those with insertions, indicating that the vast majority of L1/SVA insertions did not affect their flanking upstream methylation.

To further validate and generalize our findings, we aggregated all polymorphic insertions and plotted the average methylation trendline based on their relative position in relationship to the insertion sites from the reference genome (Supplemental Fig. S4a). We found a subtle increase of methylation flanking the polymorphic L1/*Alu* insertions, with the highest increase right at the insertion site (3.1% for L1 and 3.2% for *Alu*). This flanking methylation difference between insertion and empty site decreased as the distance to insertion position increased, and the methylation difference diminished at ∼1kb away from the insertion site (Supplemental Fig. S4a). The exact same pattern was recapitulated by both the ONT methylation data (phased ONT reads) and the WGBS data (WGBS reads of homozygous insertions to the individual’s own genome assemblies) from the five HPRC individuals (Supplemental Fig. S4b,c) (Methods). Taken together, these results suggest that polymorphic *Alu* and L1 insertions are associated with slightly elevated DNA methylation levels at their insertion sites with highly localized and small effect size.

### The exceptions: TE insertions associated with flanking region methylation change

Despite most TE insertions not affecting the methylation of their insertion sites, we also found numerous TE insertions associated with flanking methylation changes. For instance, the hypermethylated L1 insertion at chr3:85527420 (average promoter methylation level 88%) was associated with an increase in methylation of the CpG site upstream of the insertion. We plotted the methylation of each CpG site of the 5’end 1000 bp of the L1 along with both flanking regions across all 32 samples in Supplemental Fig. S5a and summarized them in Fig. 5a. In samples without this insertion, the upstream CpG site was mostly hypomethylated (average methylation 24.3%), whereas in samples with the insertion, it was generally intermediately methylated (average methylation 42.9%) (Fig. 5a). Conversely, the L1 insertion with a hypomethylated promoter (average methylation level 46.3%) at chr2:108834214 was associated with a reduction in methylation of its upstream region (from 94.2% to 74.7%) (Fig. 5b and Supplemental Fig. S5b).

**Figure 5.**
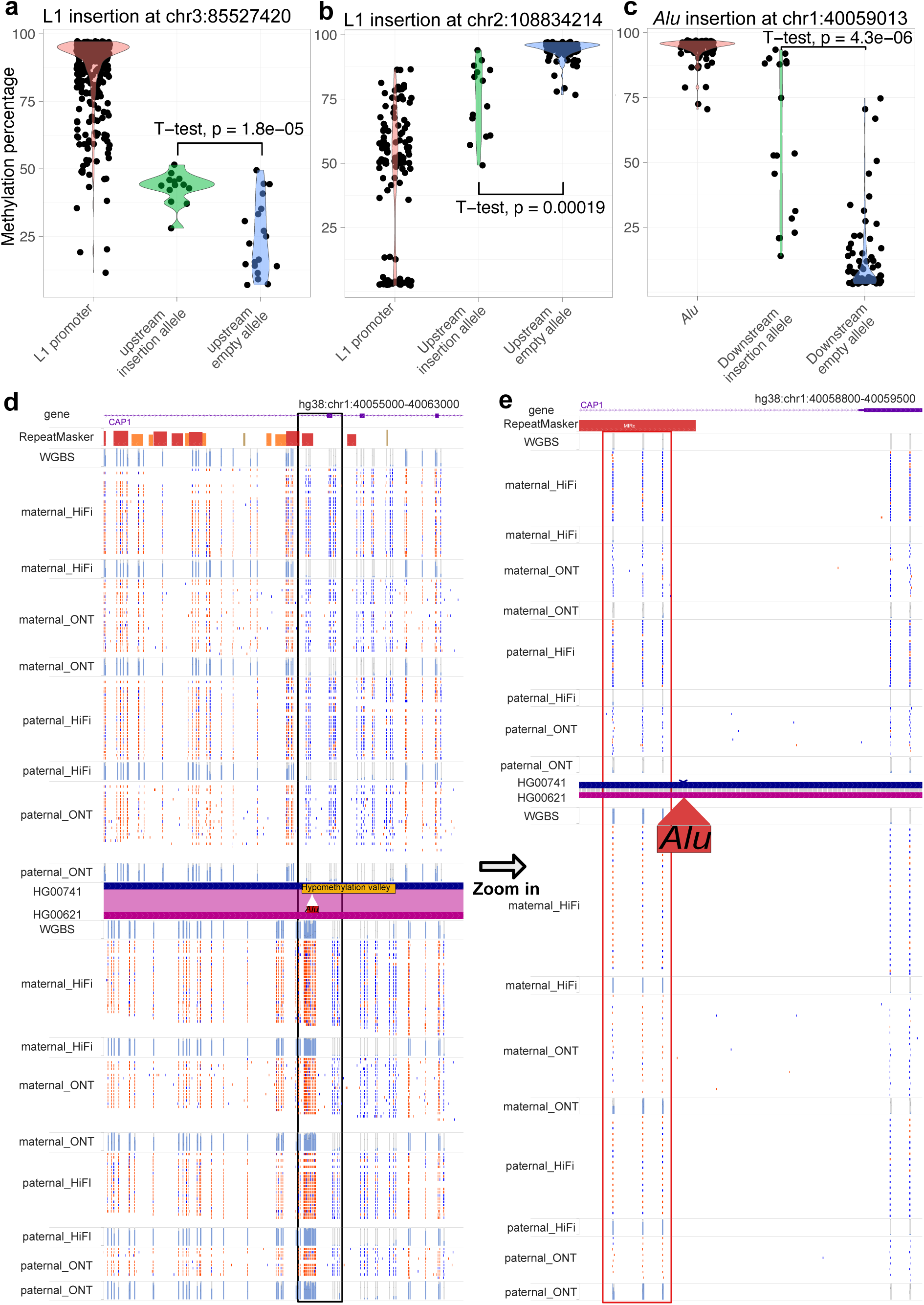
**a**: The violin plot illustrated the methylation levels of the L1 promoter at chr3:85527420. It compared the upstream CpG methylation of the insertion allele and the empty allele. Each dot represented the methylation percentage of a specific CpG site in a particular sample. A t-test was conducted to compare the matching upstream methylation levels, and the p-value is indicated on the figure. **b**: The violin plot of L1 promoter methylation at chr2:108834214 is similar to Fig.5a. The upstream CpG methylation of the insertion allele and the upstream methylation of the empty allele were shown. Each dot represented the methylation percentage of a specific CpG site in a particular sample. A t-test was performed and labeled accordingly. **c**: The violin plot of *Alu* methylation at chr1:40059013 is similar to Fig. 5a. The downstream CpG methylation of insertion allele and the downstream methylation of empty allele were shown. Each dot represented the methylation percentage of a specific CpG site in a particular sample. A t-test was performed and labeled accordingly. **d**: The WashU Epigenome Browser view of a hypermethylated *Alu* insertion within a hypomethylated valley that was present in HG00621 but absent in HG00741 characterized by WGBS, HiFi, and ONT methylation. Long-reads separated by their haploid type displayed the methylation percentage with the blue-red (low-high) two color scale, and all the methylC tracks displayed methylation percentages ranging from 0% to 100%. **e**: Zoom-in view of the methylation of the same *Alu* element from Fig. 5d. The boxed 3 CpG sites within 200bp of the insertion site underwent a change in methylation status, transitioning from hypomethylation to hypermethylation in association with the *Alu* insertion.

The impact of TE insertions on nearby genomic sequence can also be appreciated by examining heterozygous insertion in the same individual, for example, the L1 insertion at chr3:85527420 in both HG00621 and HG00741 (present only in the paternal haplotype). Both long-read methods identified the CpG site 46bp upstream of the L1 insertion was hypermethylated (average 81%) but remained hypomethylated in the remaining eight haplotypes without the insertion (average 21%) (Supplemental Fig. S5h). Interestingly, this specific polymorphic L1 was reported by Lanciano et al. to increase upstream CpG methylation in the MCF7 cell line, but was hypomethylated and had reduced proximal methylation in the 2102Ep cell line (Lanciano et al. 2024). Taken together, these results suggest that this specific polymorphic L1 insertion adopts cell type-specific DNA methylation which may spread to its immediate region upstream.

We confirmed the impact of multiple L1 loci, including the insertion at chr3:85527420 we described above. However, there were also instances where the relationship between methylation differences and L1 insertions was less evident. For instance, the methylation of the empty allele at CpG site chr14:24523788, upstream of the L1 insertion at chr14:24523704, exhibited significant variability across the HPRC samples. Meanwhile, the same CpG site was intermediately methylated among samples with the insertion allele, and the difference was not statistically significant (p=0.42) (Supplemental Fig. S5c,d). At CpG site chr5:137679380, upstream of the L1 insertion at chr5:137679083, while the methylation difference between the insertion and empty alleles was significant (p=0.0056), the high methylation variation of the empty allele made it difficult to conclude that the insertion allele methylation was influenced by the L1 insertion (Supplemental Fig. S5e,f).

Our study identified thousands of *Alu* insertions and most of them were hypermethylated. In some cases, their methylation appeared to be influenced by their genomic context (Fig. 3). Can *Alu* insertions influence the methylation of its nearby regions? We found that while the large majority of *Alu* insertions seemed to have no impact on nearby DNA methylation, exceptions did exist. For example, polymorphic *Alu* insertion at chr1:40059013 was hypermethylated (average methylation 94%), and so were 3 downstream CpG sites within 200bp (average methylation 58.1%). In samples without the *Alu* insertion, these three CpG sites were hypomethylated (average methylation 12.2%) (Fig. 5c and Supplemental Fig. S5g). A closeup view revealed that this *Alu* insertion was homozygous in HG00621 but absent in HG00741. This *Alu* was inserted within a hypomethylation valley (chr1:40058800-40060800, methylation level 4%). In HG00621, the *Alu* insertion was hypermethylated (methylation level 85%), and methylation level of the hypomethylation valley was elevated to 21% (Supplemental Fig. S5h) (Li et al. 2022; Zhuo et al. 2023). The three CpG sites 200bp 3’ to the *Alu* insertion site were lowly methylated in HG00741 (18%, 5%, 4%) but highly methylated in HG00621 (89%, 88%, 93%). On the other hand, multiple CpG sites more than 300bp downstream from the insertion site seemed to have remained their hypomethylated state (3 CpGs, 3% in HG00741, 5% in HG00621) (Supplemental Fig. S5h).

## DISCUSSION

Characterizing cytosine methylation is important for both evolutionary analysis and tissue-specific gene regulation. However, conventional bisulfite or enzyme conversion-based methods do not readily characterize methylation of repetitive regions or non-reference regions. Using 3rd generation sequencing, we can call methylation during the base calling step, greatly simplifying the methylation calling process and allowing new investigations into genomic DNA methylation. Here we compared the methylation calling results between HiFi, ONT, and WGBS from five HPRC LCL samples, and we found both HiFi and ONT can measure CpG methylation in human genome with high correlation and concordance in comparison with WGBS.

We characterized the methylation profile of non-reference TE insertions. Surprisingly, we found that the VNTR regions of non-reference SVA insertions were methylated at a lower level compared to their counterparts in the reference genome. Given the variable length of VNTR, a more specific analysis is necessary to determine their methylation levels accurately.

How are genomic insertions, in particular transposable element insertions, methylated, and how do they affect the methylation level of their flanking regions, have been a long-standing question in the field. It has promoted several paradigms, especially in the context of spreading of epigenetic modifications. For example, the hypermethylation of a B2 element close to the Aprt promoter in mouse genome can spread to nearby CpG sites until stopped by SP1 binding (Turker 2002). By examining CpG density flanking *Alu* insertions, Zhou et al. found evidence of CpG methylation spreading from old TEs to flanking regions during evolution (Zhou et al. 2020). Recently, Lanciano et al. described examples of L1 insertion in cancer cell lines affecting the methylation level of nearby regions up to 300bp upstream (Lanciano et al. 2024). It was also reported that hypomethylated L1 insertions can create hypomethylated “slopping shore” (Grandi et al. 2015). These previous studies provided enormous insights on how newly inserted transposable elements are epigenetically modified and how the modifications influence genomic neighborhood. However, most studies represented specific examples or anecdotes, leaving the question open as to what degree these examples represent the genome-wide rules or exceptions.

Taking advantage of both SV and methylation calling using the HiFi reads from 32 samples of HPRC Release1 dataset, we addressed this question by analyzing differential methylation between alleles with polymorphic insertions at a genome-wide scale. Our analysis suggests that the methylation of non-TE insertions is strongly associated with their genomic context. Consistent with Lanciano et al., we find some methylation spreading from L1 to nearby regions. However, the effect is very small (on average ∼3% increase of methylation within 300bp). The minimum spreading of methylation from polymorphic TE insertions to their flanking regions we found suggests that methylation spreading is a slow evolutionary process with limited effect after TE insertion but can potentially have large impact at an evolutionary timescale (Zhou et al. 2020).

However, our analysis was conducted on EBV-derived lymphoblastoid cell lines, which are commonly used for genome sequencing. It is crucial to note that the genomic methylation status observed in this cell line does not reflect the diverse epigenomic characteristics of various human cell types (Roadmap Epigenomics Consortium et al. 2015). Therefore, careful characterization of other cell types is still necessary to validate our findings in this cell line and to gain a deeper understanding of the dynamic methylation patterns in humans.

Because our strategy was conceptually similar to that from Lanciano et al., we directly examined the potentially polymorphic L1 insertions characterized by their study, which were reported to exhibit short-range epivariation in almost half of the cases. We found 21 out of the 87 L1 insertions from their study had sufficient HiFi coverage of both empty and insertion alleles in the HPRC dataset. The 300bp upstream methylation of empty and insertion alleles were highly correlated with R^2^ = 0.97 if they were classified as “not influenced”; with R^2^ = 0.86 if they were classified as “inconclusive”; and with R^2^ = 0.61 if they were classified as “influenced” (Supplemental Fig. S6), as defined in Lanciano et al. (Lanciano et al. 2024). Thus, those L1 insertions that were determined by Lanciano et al. to exhibit short-range epivariation were indeed associated with slightly increased differences between alleles with the insertion and the empty alleles, replicated in our own dataset.

However, the empty allele and insertion allele from the same locus were still strongly correlated with each other, indicating the effect size of L1 influence was small. In contrast, our HPRC dataset contained a much larger sample size, and we determined that the flanking methylation difference between the insertion allele and the empty allele was less than 25% for most cases (287 out of 292 L1 insertions and 4519 out of 4693 *Alu* insertions) (Fig. 4). Thus, we concluded that the majority of polymorphic TE insertions do not change their flanking region DNA methylation, while there exist numerous exceptions with a limited short-range influence.

To visualize the methylation of non-reference insertions, we mapped reads to the high-quality individual assemblies and displayed them on the WashU Epigenome Browser using the genome comparison feature (Li et al. 2022; Zhuo et al. 2023). However, this approach is limited by the underlying pairwise alignment and more crucially, the availability of high-quality individual assemblies. We hope the development of pangenome based tools would facilitate data display of non-reference sequences in the future.

With the ongoing effort and continuing release of high-quality data from HPRC, we envision the publicly available long read methylation data can be used to address other questions in the biomedical field, and we invite other scientists to explore the rich, high-fidelity HPRC dataset (Liao et al. 2023).

## METHODS

### Genome assembly and scaffolding

The genome assemblies used in this study were produced for the draft human pangenome, combining PacBio HiFi long-read and Illumina short-read sequencing for de novo assembly with parental short-read data used to for phasing of these assemblies. Phased contig assembly was performed using hifiasm (Cheng et al. 2021). Post-assembly, the RagTag toolset was employed for reference-guided scaffolding, organizing contigs into structured, phased chromosomal assemblies (Alonge et al. 2022). Validation was performed using BioNano optical mapping, in order to detect and amend potential assembly inaccuracies (Lam et al. 2012). For each of the 10 assemblies derived from the five diploid samples, we created a whole-genome pairewise alignment BAM file with hg38 using minimap2 (Li 2018).

### Whole Genome Bisulfite Sequencing (WGBS)

The cytosine methylation was estimated by mapping to reference genome hg38 or the individual assemblies using the standard WGBS pipeline Bismark as described by Lee et al. (Krueger and Andrews 2011; Lee et al. 2020a).

### PacBio HiFi Methylation

PacBio single-molecule high fidelity (HiFi) Circular Consensus Sequencing (CCS) reads with kinetics tags can be generated using pbccs (https://github.com/nlhepler/pbccs). We then predicted cytosine methylation probability per molecule in all CpG context by applying primrose to PacBio HiFi reads (https://github.com/PacificBiosciences/jasmine). The CpG methylation probability was saved as the ML and MM tags in the BAM file following standard BAM format convention (http://samtools.github.io/hts-specs/SAMtags.pdf). Unaligned HiFi BAM files were aligned to human reference genome hg38 using PacBio pbmm2 (https://github.com/PacificBiosciences/pbmm2), and CpG methylation percentage on the reference genome were calculated using PacBio pb-CpG-tools (https://github.com/PacificBiosciences/pb-CpG-tools). We ran the methylation percentage calculation using default model “PB-model”.

### Nanopore ONT methylation

We downloaded raw sequencing signal of ONT Nanopore (FAST5 files) of the same five samples and performed methylation calling using basecaller guppy with methylation model remora (https://github.com/nanoporetech/remora). CpG methylation probability of ONT reads were estimated using Nanopore basecaller Guppy v6 with remora methylation model (v9.4.1) (https://community.nanoporetech.com). The CpG methylation probability were saved as the standard ML and MM tags in the BAM format (http://samtools.github.io/hts-specs/SAMtags.pdf). ONT unaligned BAM files were aligned to human reference genome hg38 using minimap2 (https://github.com/lh3/minimap2), and methylation percentage were calculated using modbam2bed (https://github.com/epi2me-labs/modbam2bed).

### Methylation comparison

With genome-wide methylation percentage of all three methods calculated, we made pairwise correlation heatmap, PCA plot and calculated the mean average error, 20% concordance, and Pearson correlation using R (Wickham 2009; Emerson et al. 2013). We also applied DSS to define pairwise differential methylation regions (DMRs) (Wu et al. 2015; Feng et al. 2014).

### Long reads phasing

We used HiPhase for HiFi reads phasing (Holt et al. 2024). To determine in each phase block which haplotype is paternal, and which is maternal, we downloaded HiFi reads used for constructing paternal and maternal assemblies from hifiasm output gfa files (HG00621, HG00741, HG01952, HG01978, HG03516). We labeled the haplotype paternal if the number of reads labeled with it from the same phase block used for paternal assembly is >10 times higher than that used for maternal assembly, otherwise maternal. Once we separated HiFi reads to paternal and maternal with whatshap (Patterson et al. 2015), we aligned them to the individual paternal and maternal assemblies, accordingly using pbmm2 (https://github.com/PacificBiosciences/pbmm2). At last, we calculated the genome-wide methylation percentage on individual assemblies using pb-CpG-tools (https://github.com/PacificBiosciences/pb-CpG-tools).

To phase ONT reads according to the individual assemblies, we created a diploid VCF file for each sample using dipcall (https://github.com/lh3/dipcall) (Li et al. 2018) and haptagged all reads using haplotype information from the VCF file. We then separated paternal and maternal reads using whatshap (Patterson et al. 2015) and aligned them to paternal and maternal assembly using minimap2 (Li 2018, 2021), respectively. At last, we converted the bam file to methylbed files with modbam2bed (https://github.com/epi2me-labs/modbam2bed).

### Individual genome visualization

For each pairwise genome alignment between hg38 and individual genome, we created a genome-align track file using the pairwise genome alignment BAM file (Zhuo et al. 2023). We built individual genome assembly of both maternal and paternal chromosomes of the five samples (10 assemblies in total) and aligned each of them to the hg38 to create genome-align tracks. We also phased HiFi reads using HiPhase using alignment to hg38 and assigned haplotypes to paternal/maternal for each phasing block based on hifiasm assembly construction graph (Holt et al. 2024; Cheng et al. 2021). We then aligned maternal and paternal HiFi reads to the maternal/paternal assembly separately for each of the five samples. Applying the genome-align track we anchored gene, repeat, and CpG island annotations from hg38 onto the individual genomes. We colored each CpG site by their methylation prediction (red: methylated; blue: unmethylated) per HiFi read using the modbed tracks and displayed piled up methylation level using the methylC track. Both modbed and methylC tracks were mapped to the individual genome. Combining genome-align, methylC, and modbed track files, we created three datahubs on the WashU Epigenome browser representing HiFi, ONT, and WGBS data (https://epigenomegateway.wustl.edu/browser2022/?genome=hg38&hub=https://wangcluster.wustl.edu/∼xzhuo/hifi_methylation/hifi.all.json, https://epigenomegateway.wustl.edu/browser2022/?genome=hg38&hub=https://wangcluster.wustl.edu/∼xzhuo/hifi_methylation/ont.all.json, https://wangcluster.wustl.edu/∼xzhuo/hifi_methylation/wgbs_individual/wgbs.all.json) (Li et al. 2022; Zhou et al. 2014; Li et al. 2023).

After aligning WGBS reads to the individual assembly as described above, we also ploted the WGBS methylation of non-reference *Alu* insertion at chr11:59565869 using Methylartist (Ewing et al. 2020; Cheetham et al. 2022).

### Structural variation and TE annotation

We obtained the structural variations of each sample from HPRC consortium (Download) (Liao et al. 2023). We merged all the called SVs from the 32 samples using bcftools merge (Danecek et al. 2021). To further annotate transposable element insertions among all structural variations, we ran MELT-LRA and PALMER2 on HiFi assemblies and xTEA on the same PacBio HiFi library mapped to hg38 (Gardner et al. 2017; Zhou et al. 2024; Chu et al. 2021). For *Alu*, L1 and SVA insertions, we included MEIs called by at least two of three methods. MELT-LRA did not call HERVK insertion, and we manually verified 25 out of 34 HERVK insertions identified by either xTEA or PALMER2. We summarized MEI caller output in supplementary (Supplemental Table S1). We then intersected the coordinates of TE insertions with SV and annotated which SVs are TE insertions (Supplemental Table S2).

We developed a script to extract non-reference insertions. Using a BED file of annotated insertion sites, it can extract inserted and soft-clipped sequences from raw reads and generate a new unmapped modBAM file of inserted sequences with base modification tags (MM and ML) (bam.extractInsertion.py). We can then align the unmapped modBAM file to their respected TE consensus (L1HS, *Alu*Y, SVA_F, LTR5_Hs and HERVK-int from dfam database) to create CpG methylation pileup results (Wheeler et al. 2012; Hubley et al. 2015).

To calculate average methylation of certain regions, we extracted methylation prediction value of each CpG site of each read from bam files with methylation tags with a python script (python bam.Mmtag.regions.oo.py -b <bam> -r <insertion.bed> -o <ins.methylC.reldist.txt> -f <bed>) available in Github repository (https://github.com/xzhuo/modbamUtil)

To evaluate the relationship between CpG methylation and indels, we aggregated methylation prediction of all 32 samples and calculated the average CpG methylation of each CpG Island (CGI) (Quinlan and Hall 2010).

### reference TE methylation

We extracted 52802 *AluY*, 2746 *AluYa5*, 36 *AluYa8*, 2111 *AluYb8*, and 301 *AluYb9* from hg38 using repeatMasker annotation. similarly, we extracted 124 SVA_E, 331 SVA_F, 481 LTR5 as reference TEs. We also downloaded 146 active L1 on hg38 from L1 Base2 (Penzkofer et al. 2017). We then take their DNA sequences from the hg38 reference and their corresponding methylation from piled up bedMethyl files and generated new modBam files to bind methylation percentage to the sequence as modified base (bam.creation.py). At last, we mapped the new BAM file to TE consensus using Minimap2 to align CpG methylation percentage to the TE consensus (L1HS, *Alu*Y, SVA_F, LTR5_Hs and HERVK-int) (Wheeler et al. 2012; Hubley et al. 2015).

Reference or non-reference insertions are relative to the reference genome chosen for alignment, and non-reference insertions can be aligned to the genome if we chose the individual assembly as the “reference”. Therefore, the sample process can be applied to extract “non-ref” insertions on the hg38 after align long reads to the individual assembly.

### Insertion and flanking methylation calculation

We selected non-reference insertions from annotated SVs and used a custom python script in the Github repository (https://github.com/xzhuo/modbamUtil) to calculate average methylation of the insertion and flanking regions (python read.methylation.mean.py -i <input> -o <output> -s <start> -l <length> -c).

### CpG island methylation and INDEL frequency calculation

Unmasked CpG island annotation file was downloaded from UCSC (https://hgdownload.soe.ucsc.edu/goldenPath/hg38/database/cpgIslandExtUnmasked.txt.gz). For each CpG site on hg38, we calculated the median methylation percentage of the 32 HiFi methylation. We then calculated the average methylation percentage of each CpG island using the median methylation.

After separating CpG islands to hypermethylated and hypomethylated ones using 50% methylation as cut-off, we calculated the INDEL frequency of different sizes within CpG islands. To create the background INDEL frequency in the genome, we shuffled CpG islands in the genome and repeated the calculation.

We also repeated the same calculation using same methylation data and INDEL callset from 1000 genome project and HGSVC project (The 1000 Genomes Project Consortium 2015; Ebert et al. 2021).

## DATA ACCESS

All 32 PacBio HiFi methylation data of HPRC year1 samples are available at https://s3-us-west-2.amazonaws.com/human-pangenomics/index.html?prefix=submissions/560047a6-6d16-4b0c-aac9-7d0c83e2188e--HIFI-METHYLATION-READS/.

The Structural variation annotation is available at https://s3-us-west-2.amazonaws.com/human-pangenomics/index.html?prefix=submissions/B581EBA7-8BDE-4C7C-9DEA-78B99A051155--Yale_HPP_Year1_Variant_Calls/.

