## Supplemental Fig. S1 for "Characterizing cytosine methylation of polymorphic human transposable element insertions using human pangenome resources"

**Supplemental Fig. S1a**

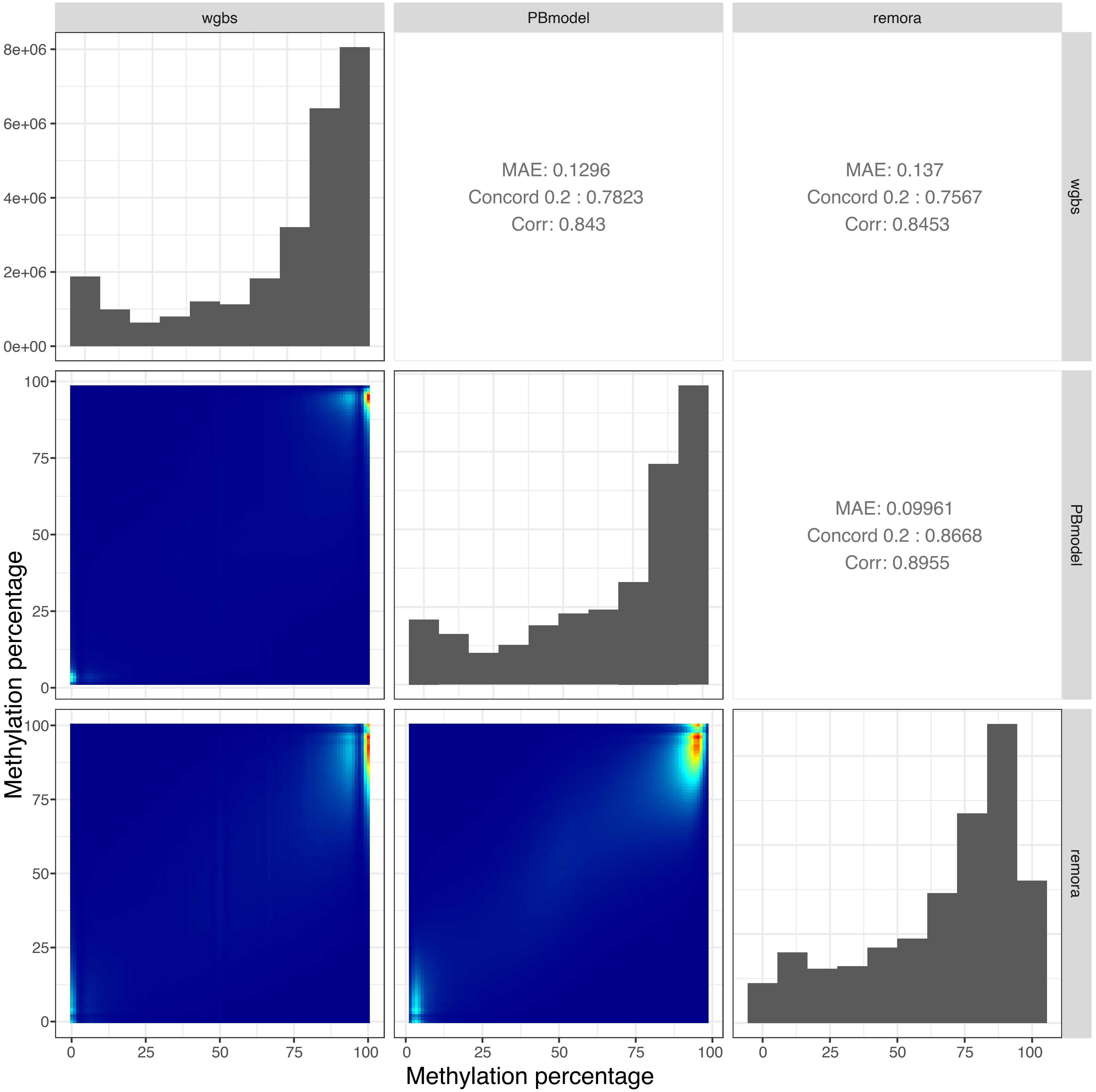

### Supplemental Fig. S1b

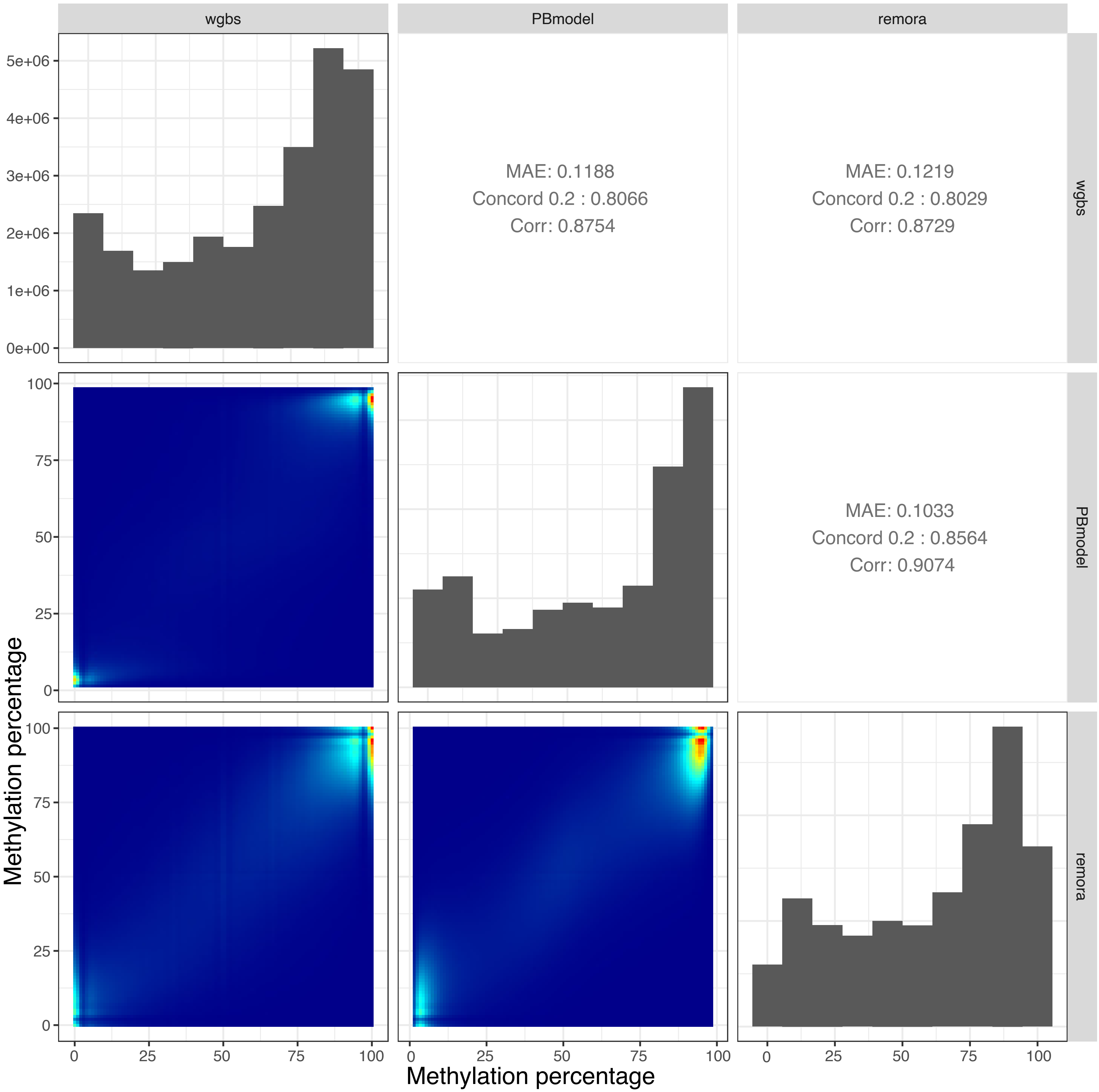

### Supplemental Fig. S1c

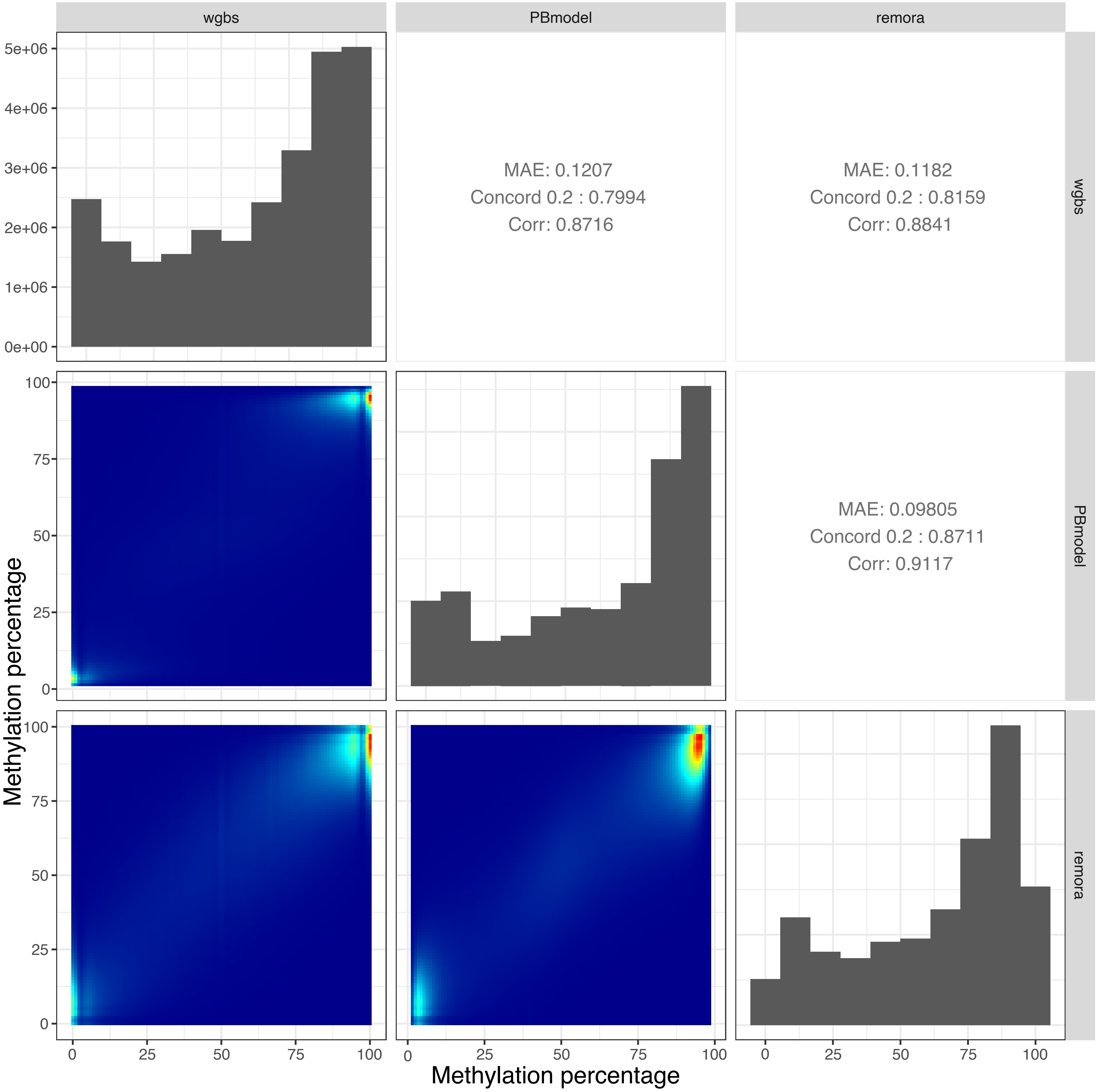

**Supplemental Fig. S1d**

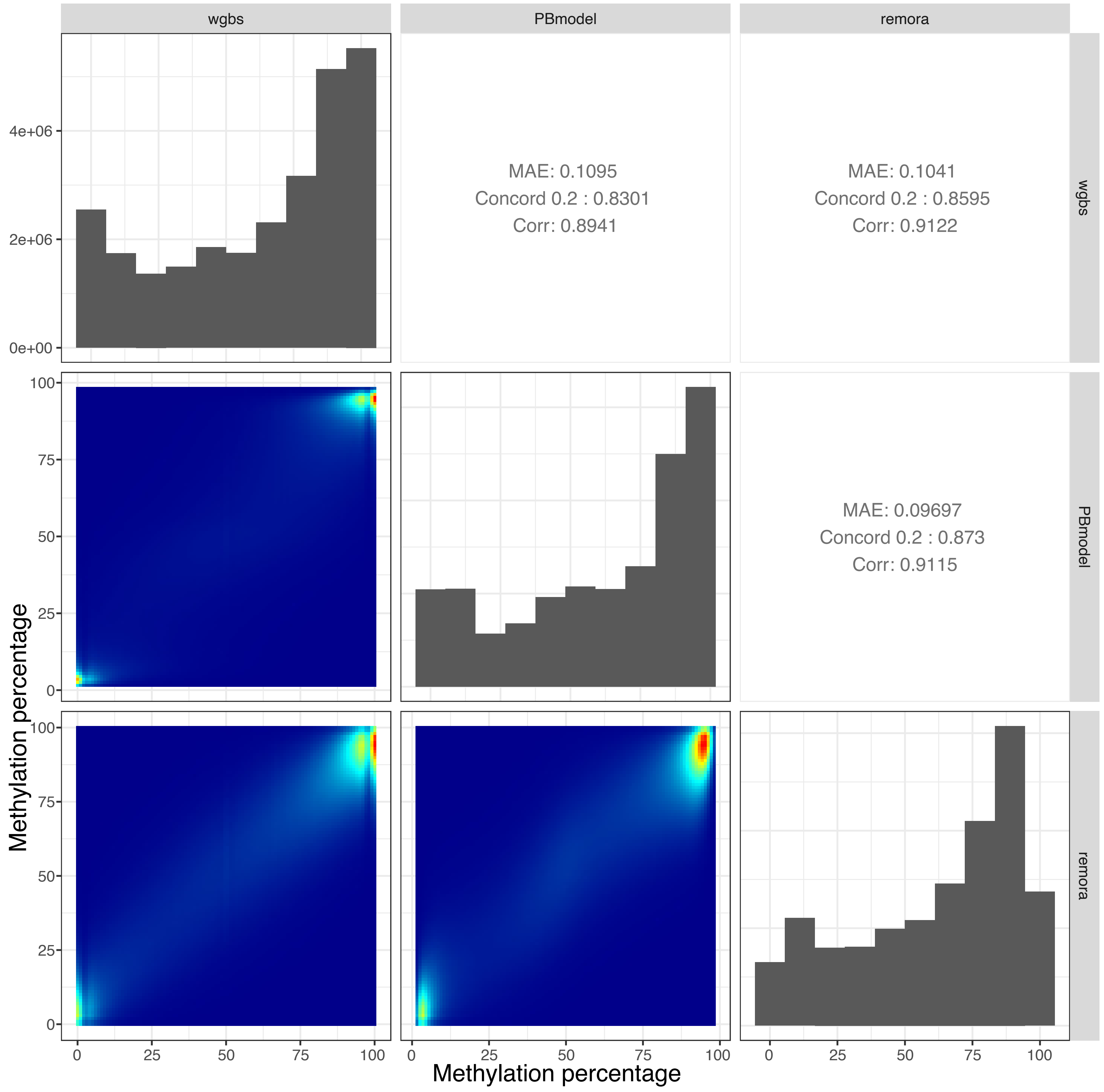

**Supplemental Fig. S1e**

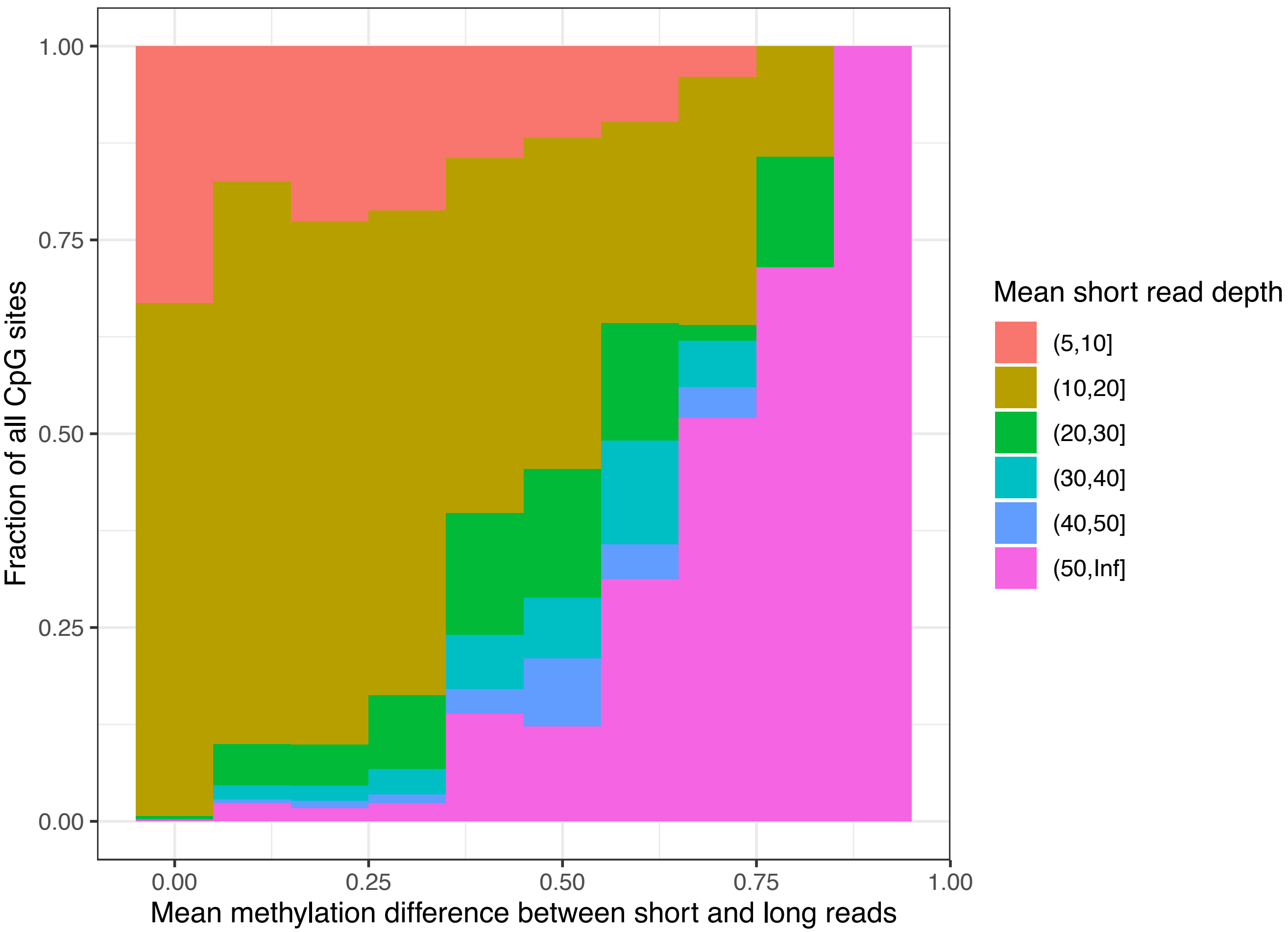

**Supplemental Fig. S1f**

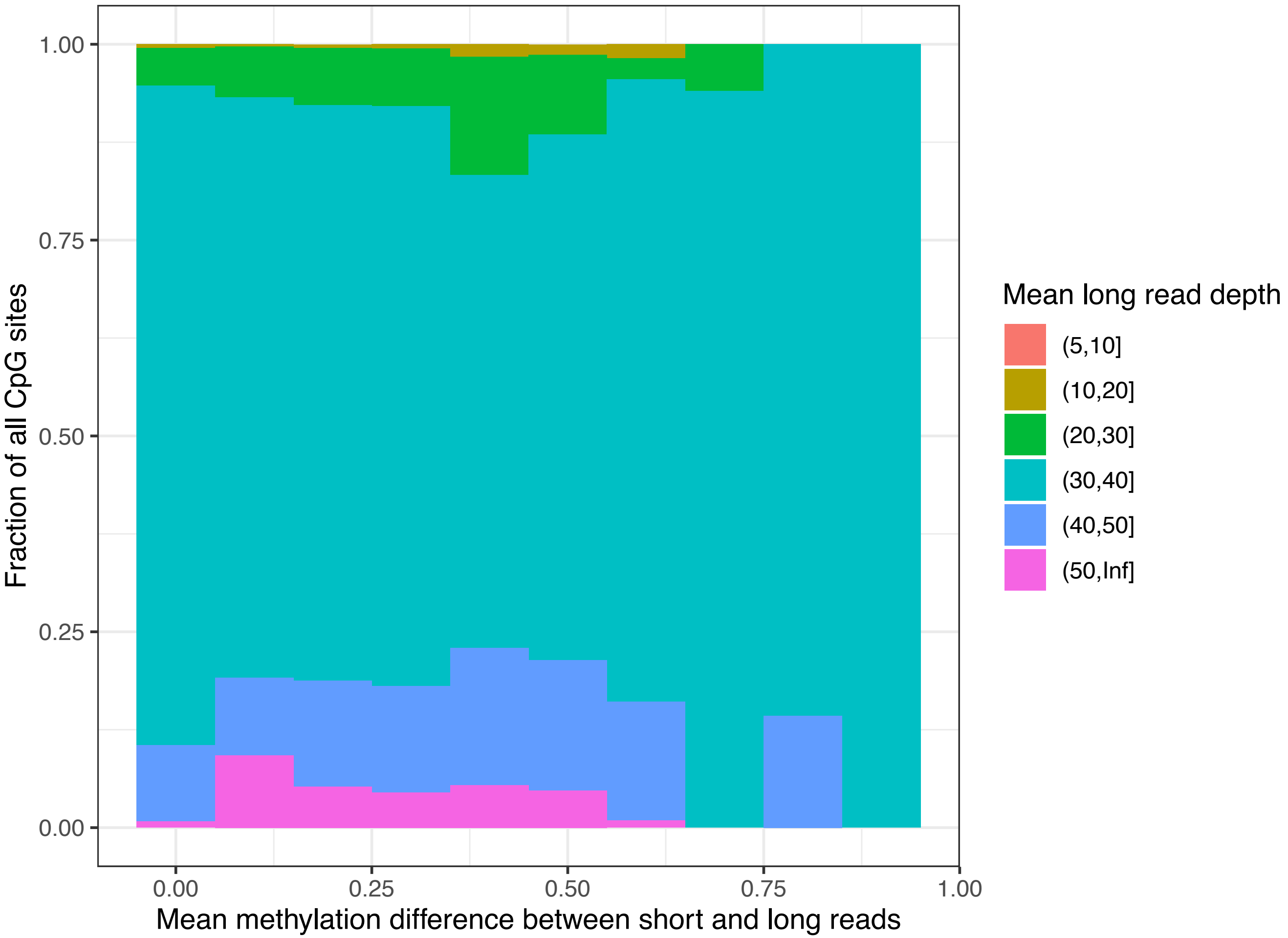

**Supplemental Fig. S1g**

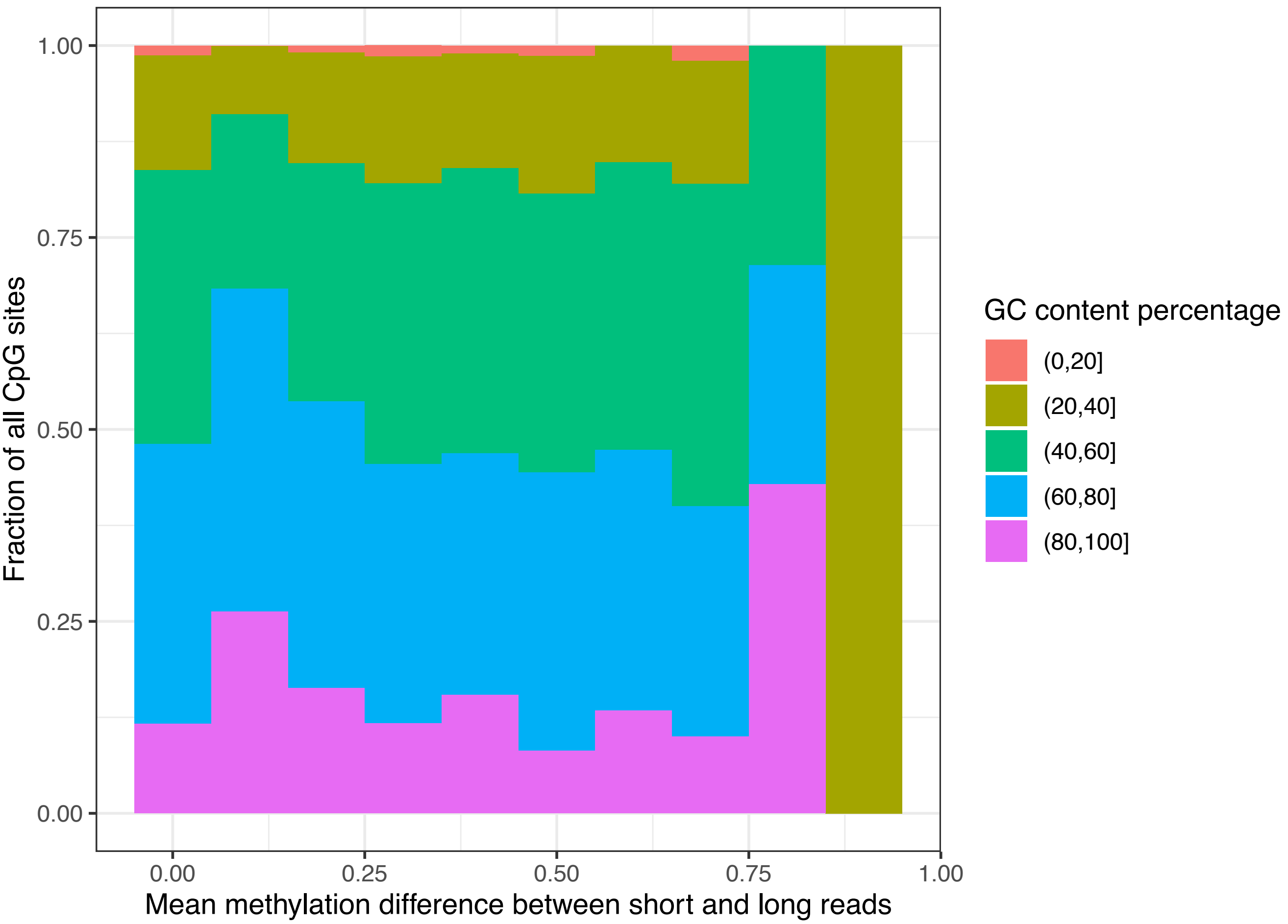

### Supplemental Fig. S1h

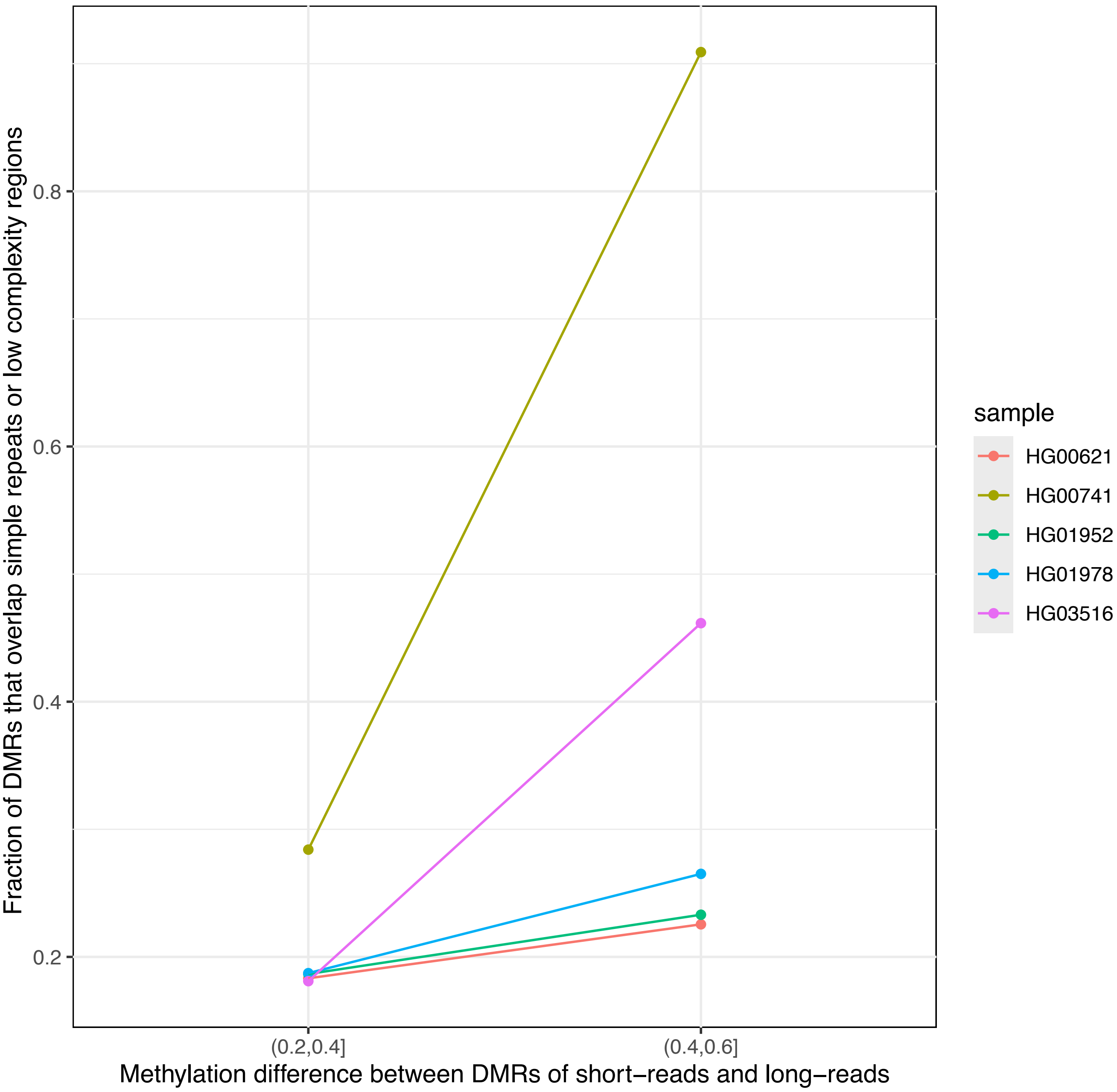
