## Supplemental Fig. S2 for "Characterizing cytosine methylation of polymorphic human transposable element insertions using human pangenome resources"

#### Supplemental Fig. S2a

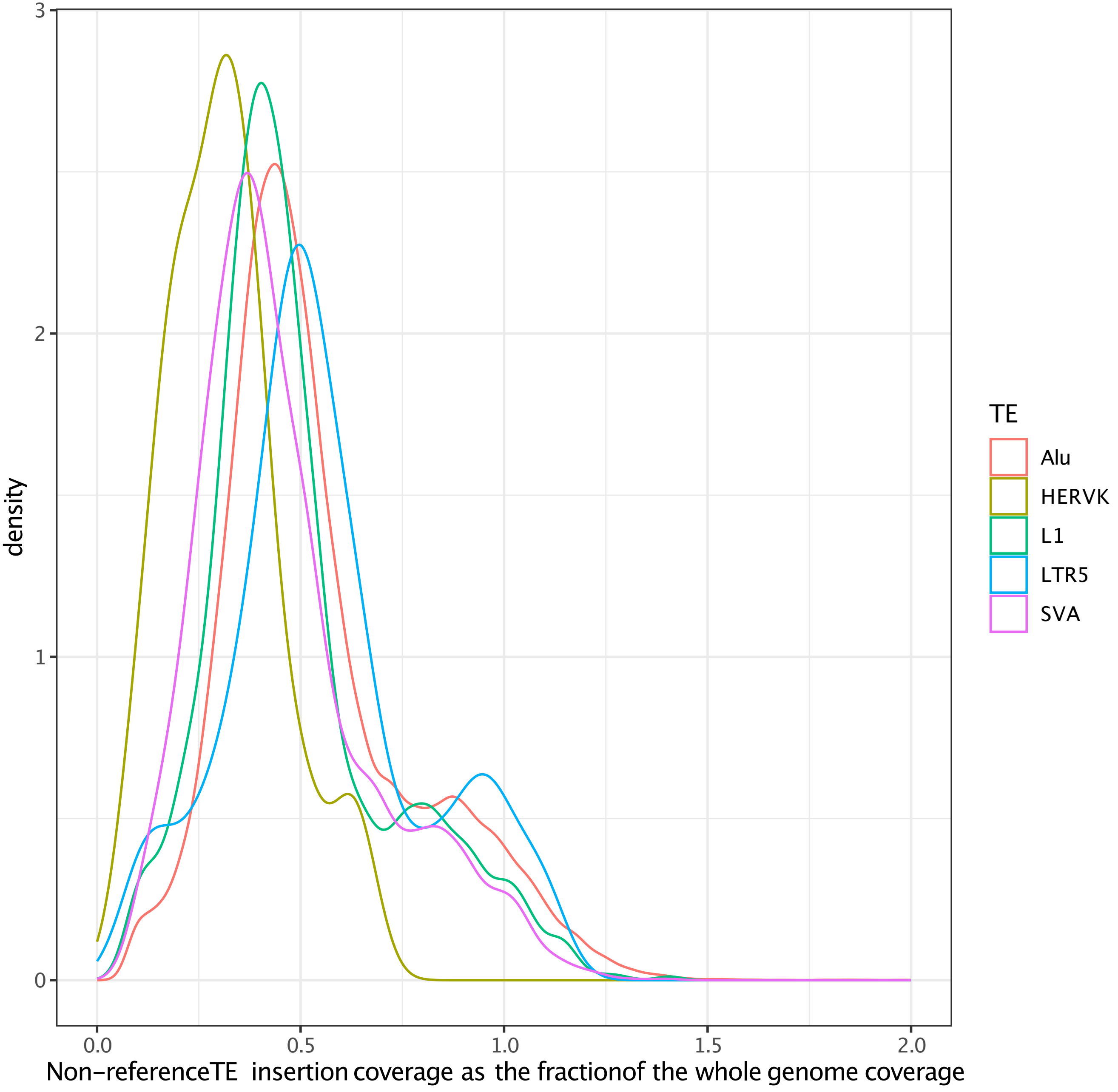

**Supplemental Fig. S2b**

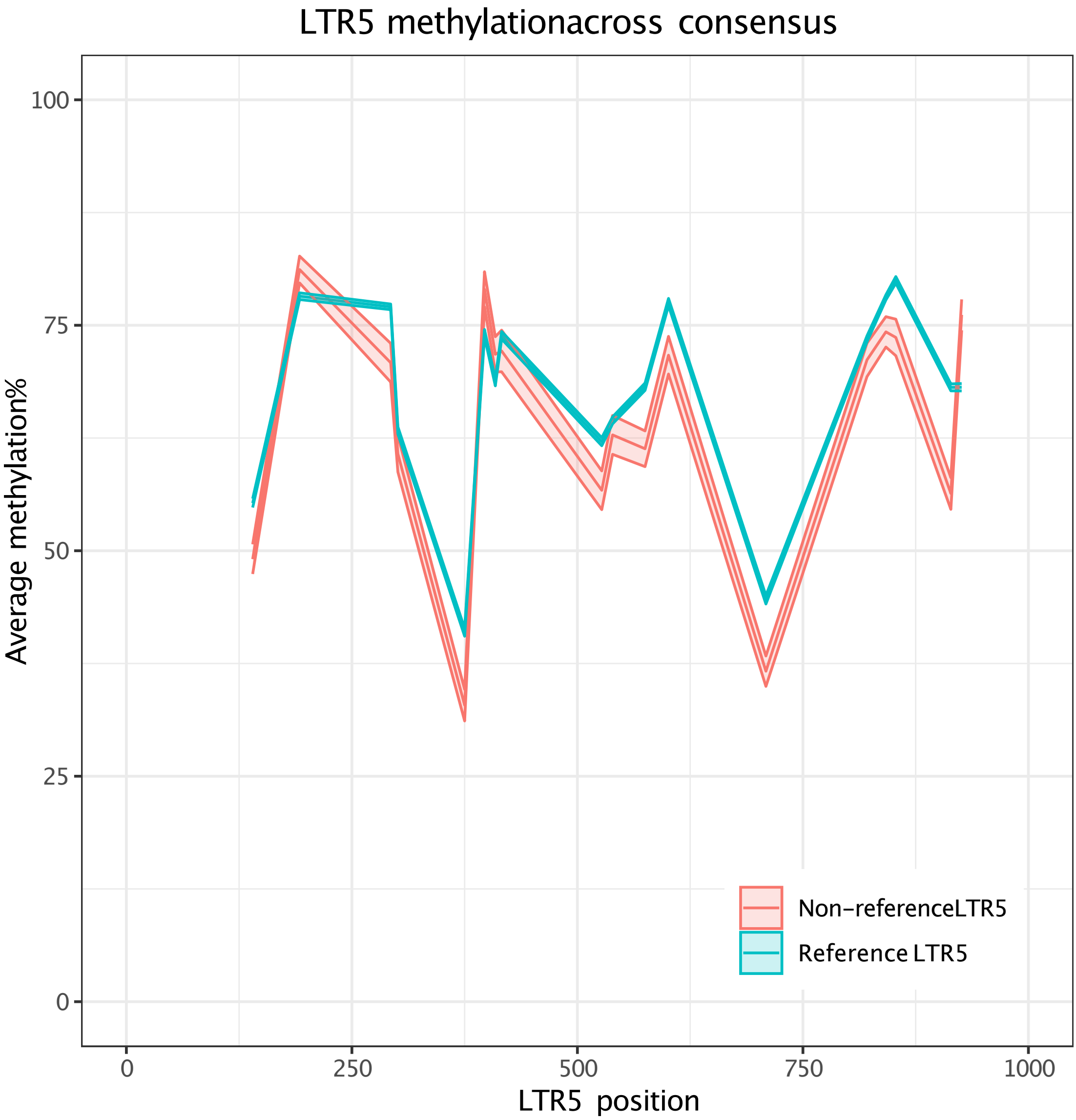

#### Supplemental Fig. S2c

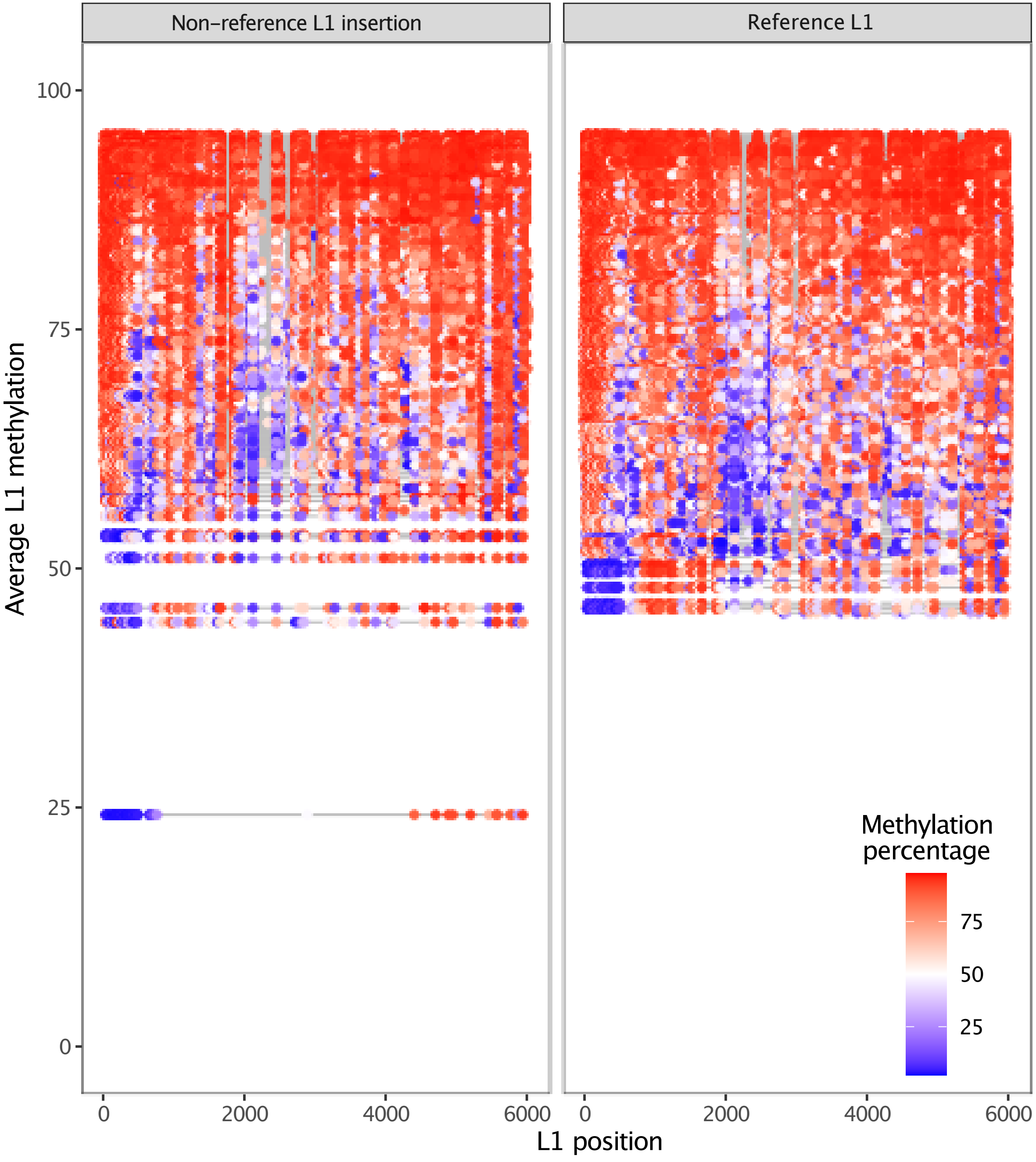

#### Supplemental Fig. S2d

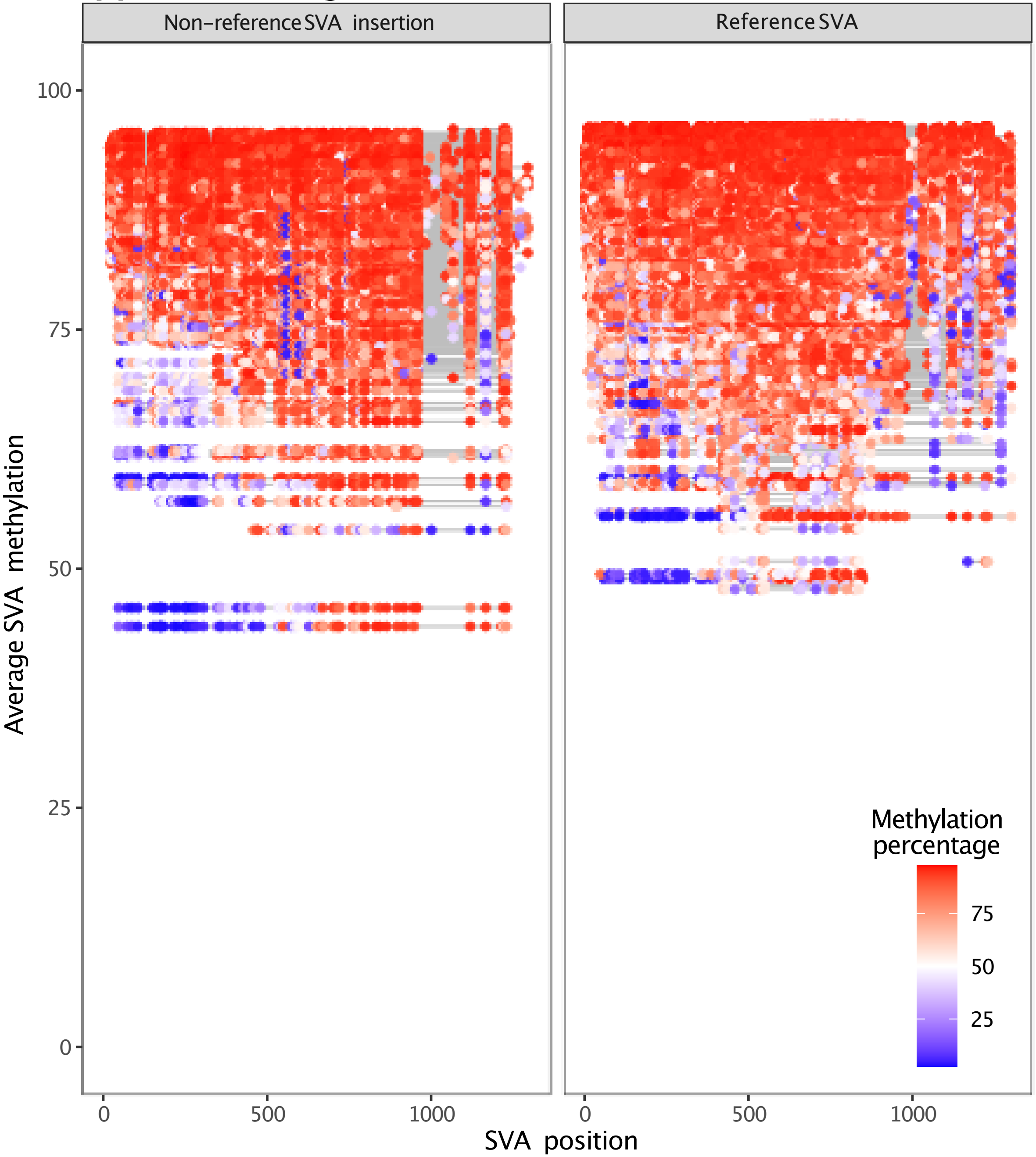

### Supplemental Fig. S2e

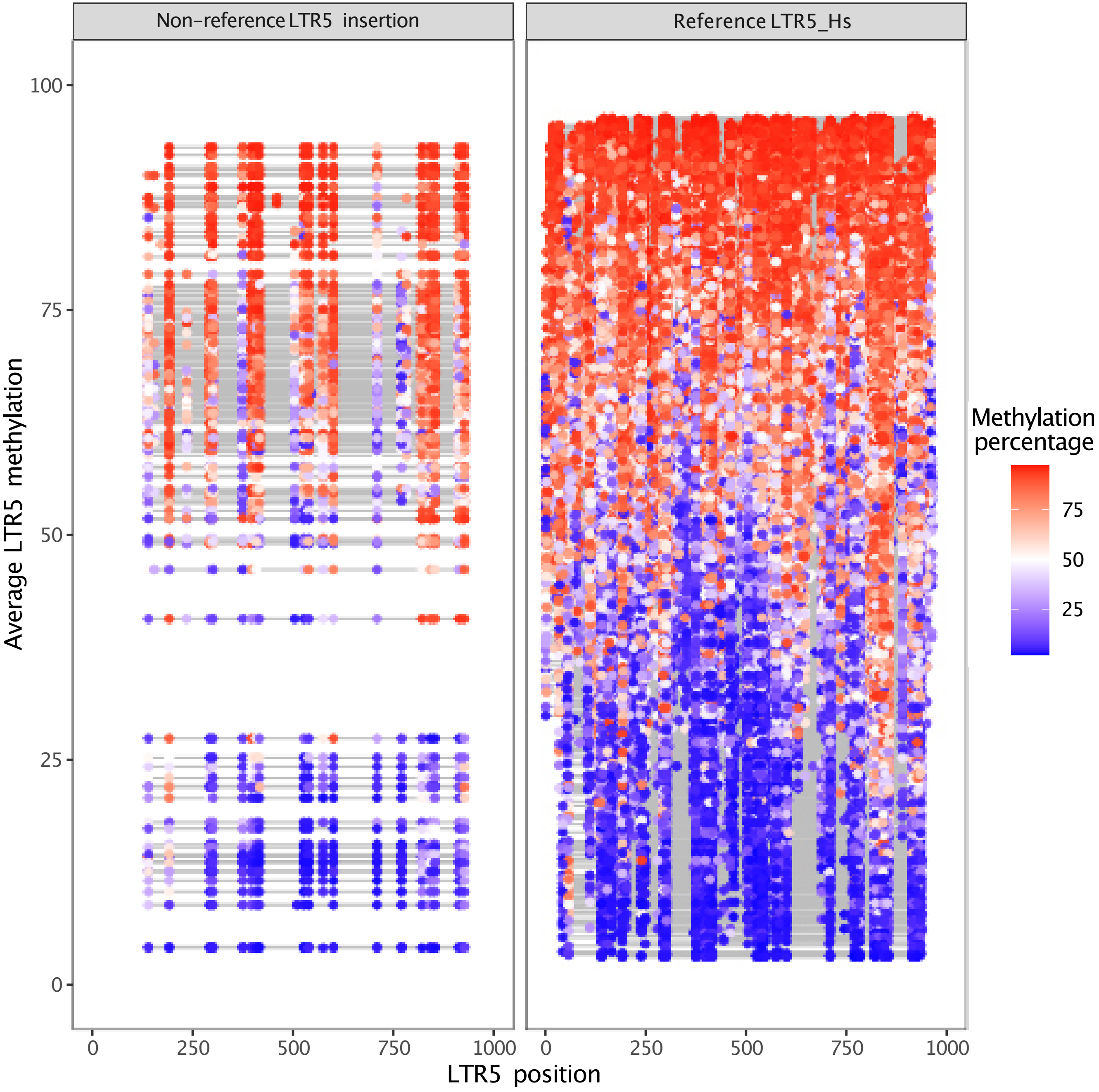

#### Supplemental Fig. S2f

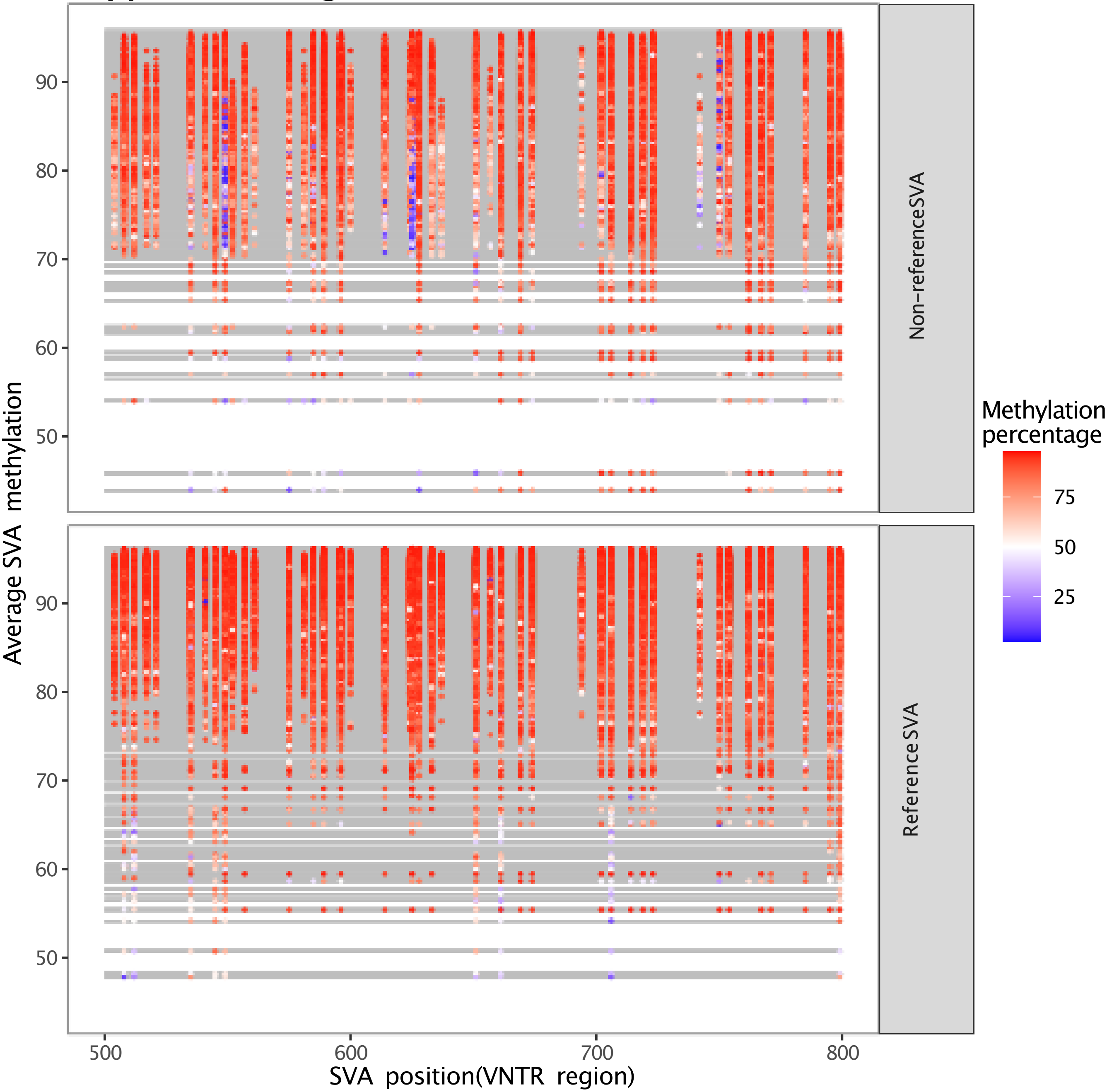

**Supplemental Fig. S2g**

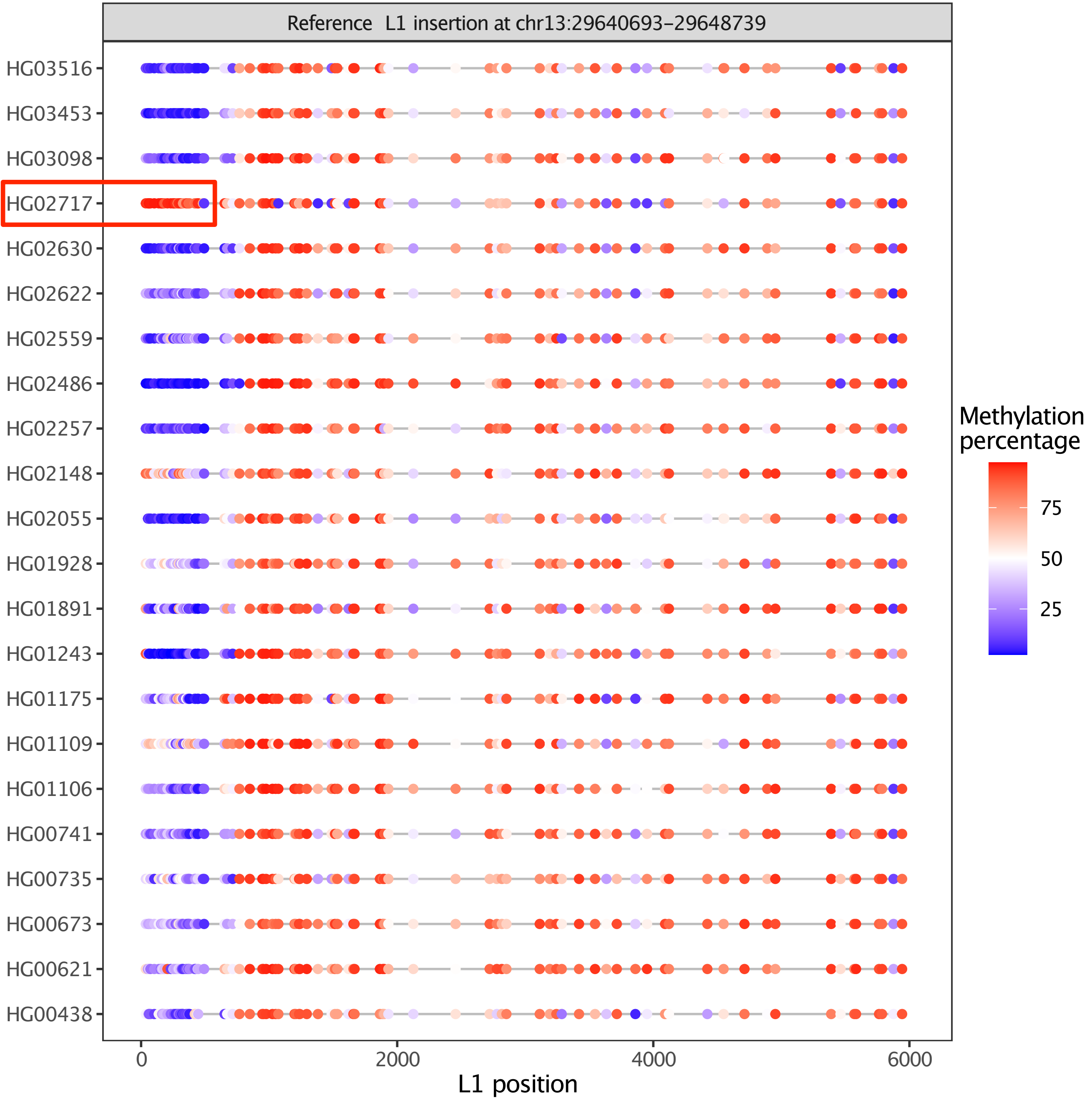
