## Supplemental Fig. S3 for "Characterizing cytosine methylation of polymorphic human transposable element insertions using human pangenome resources"

### Supplemental Fig. S3a

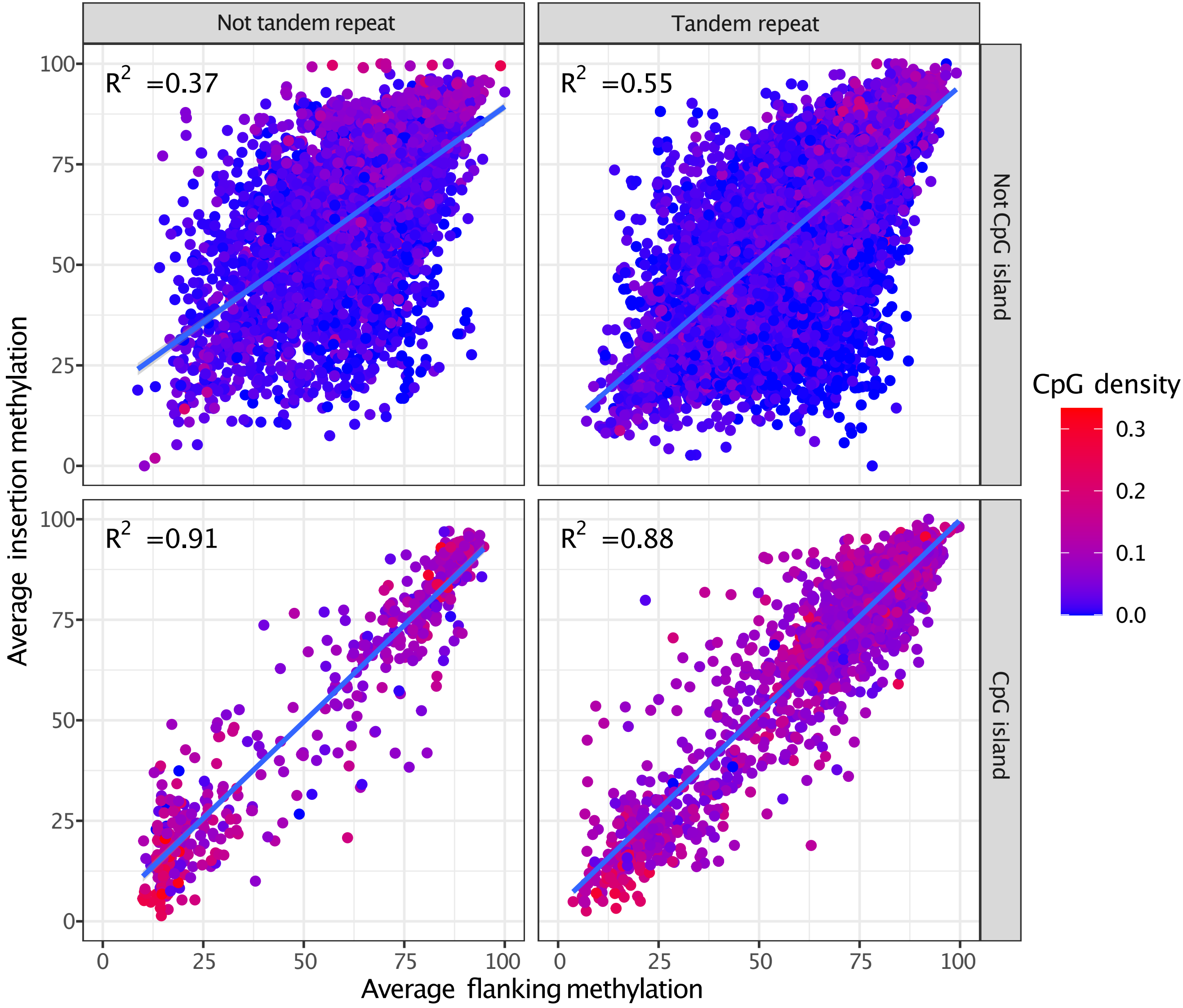

### Supplemental Fig. S3b

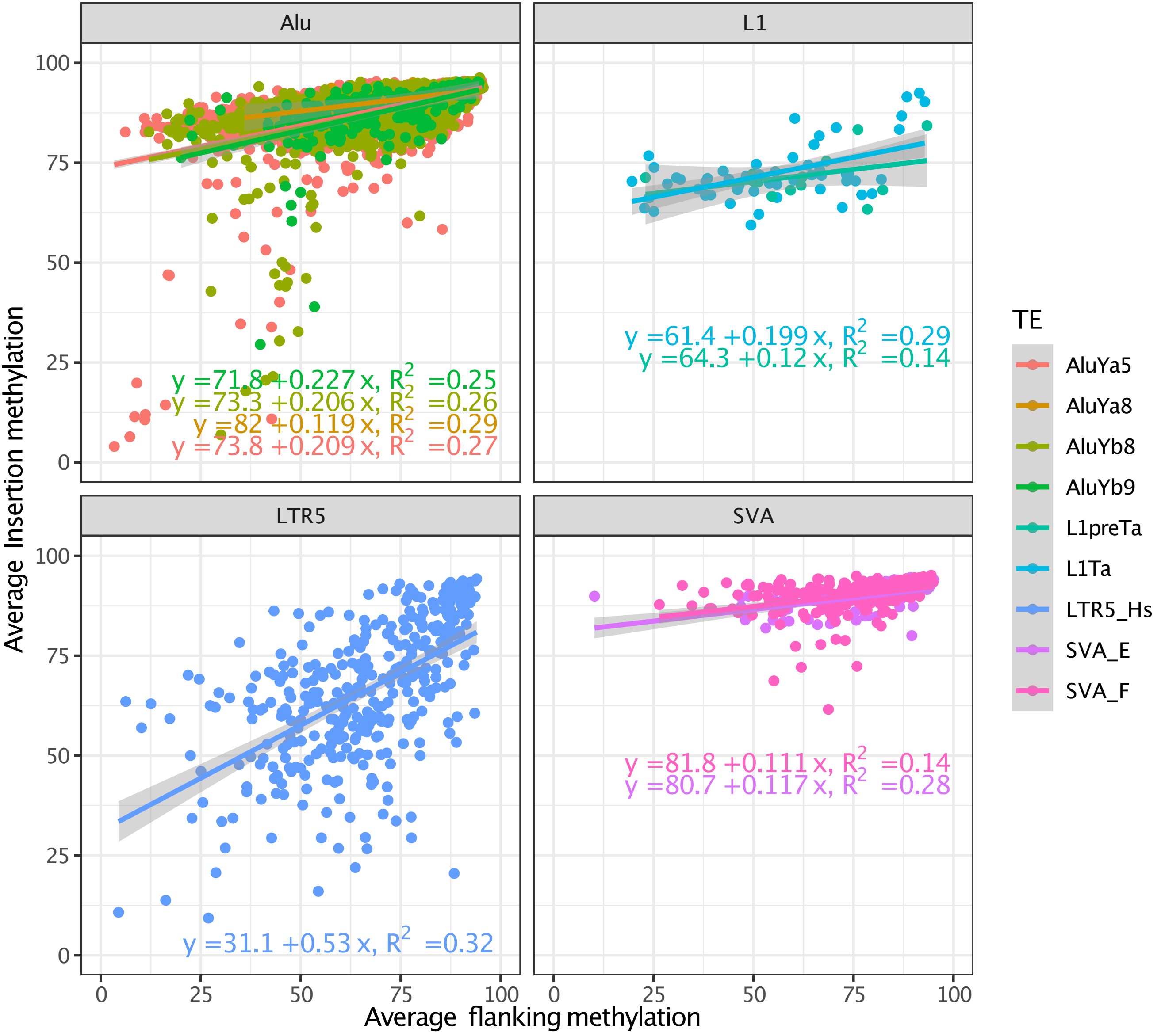

### Supplemental Fig. S3c

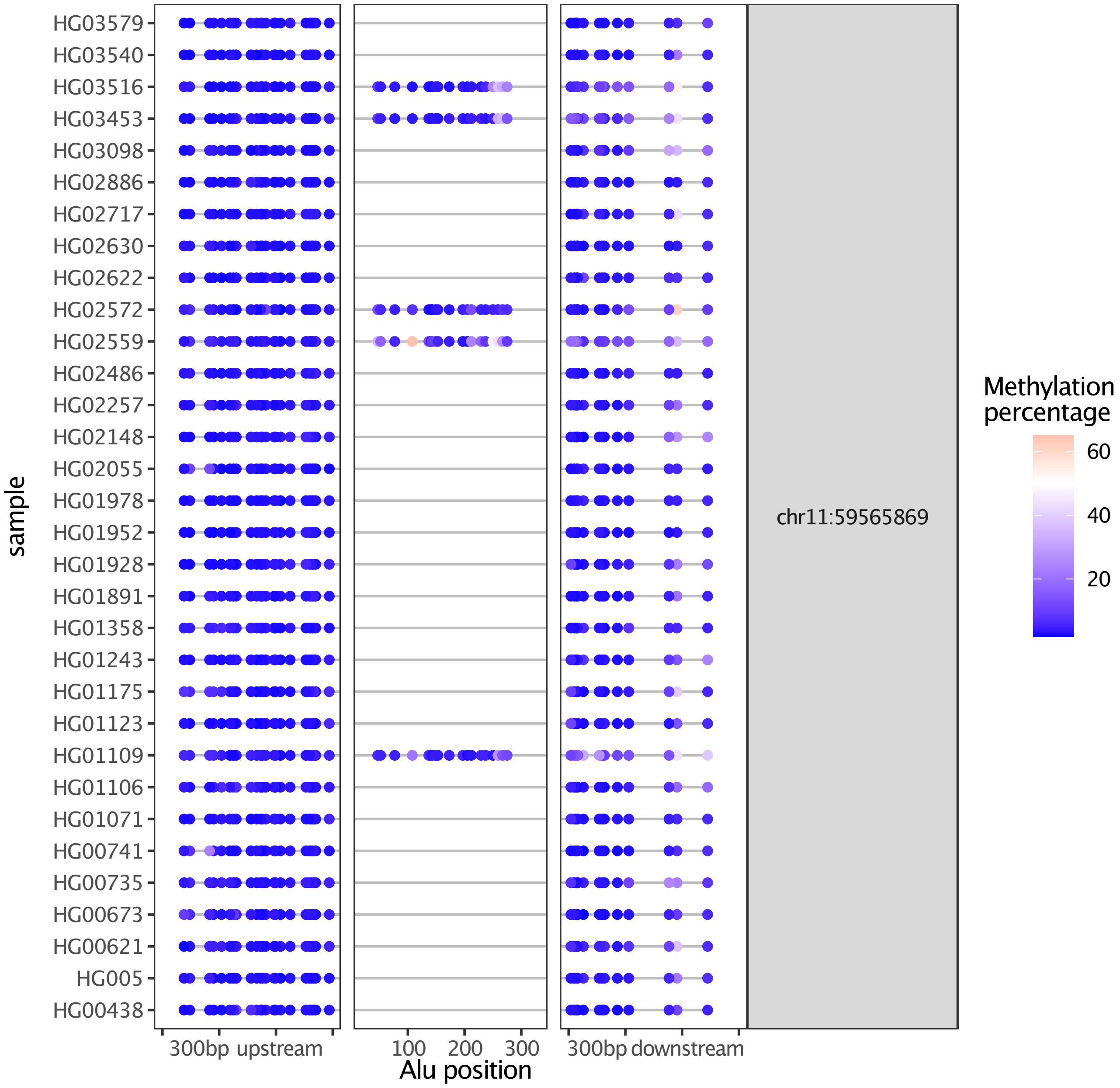

### Extended Data Figure 3d

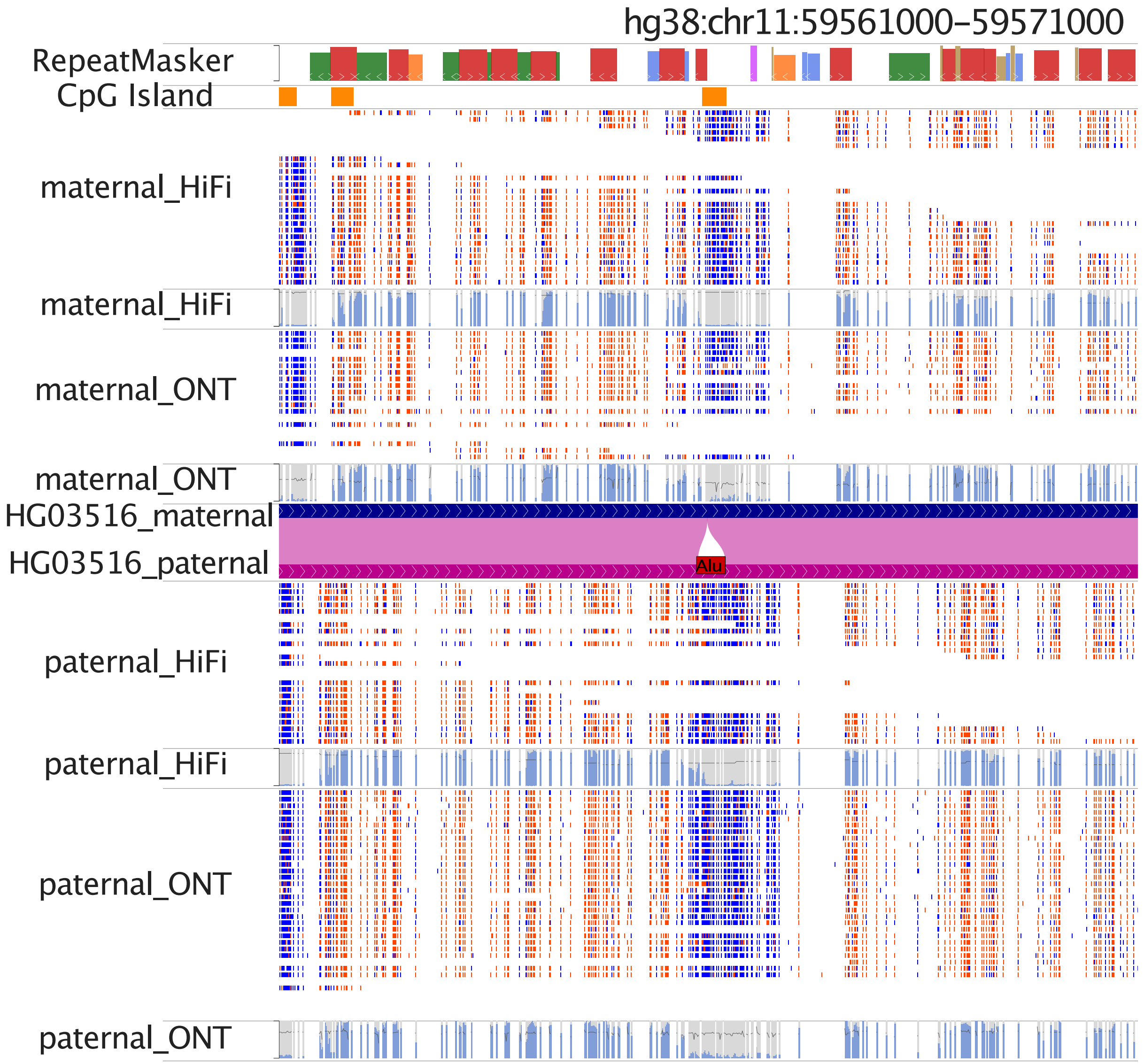

### Supplemental Fig. S3e

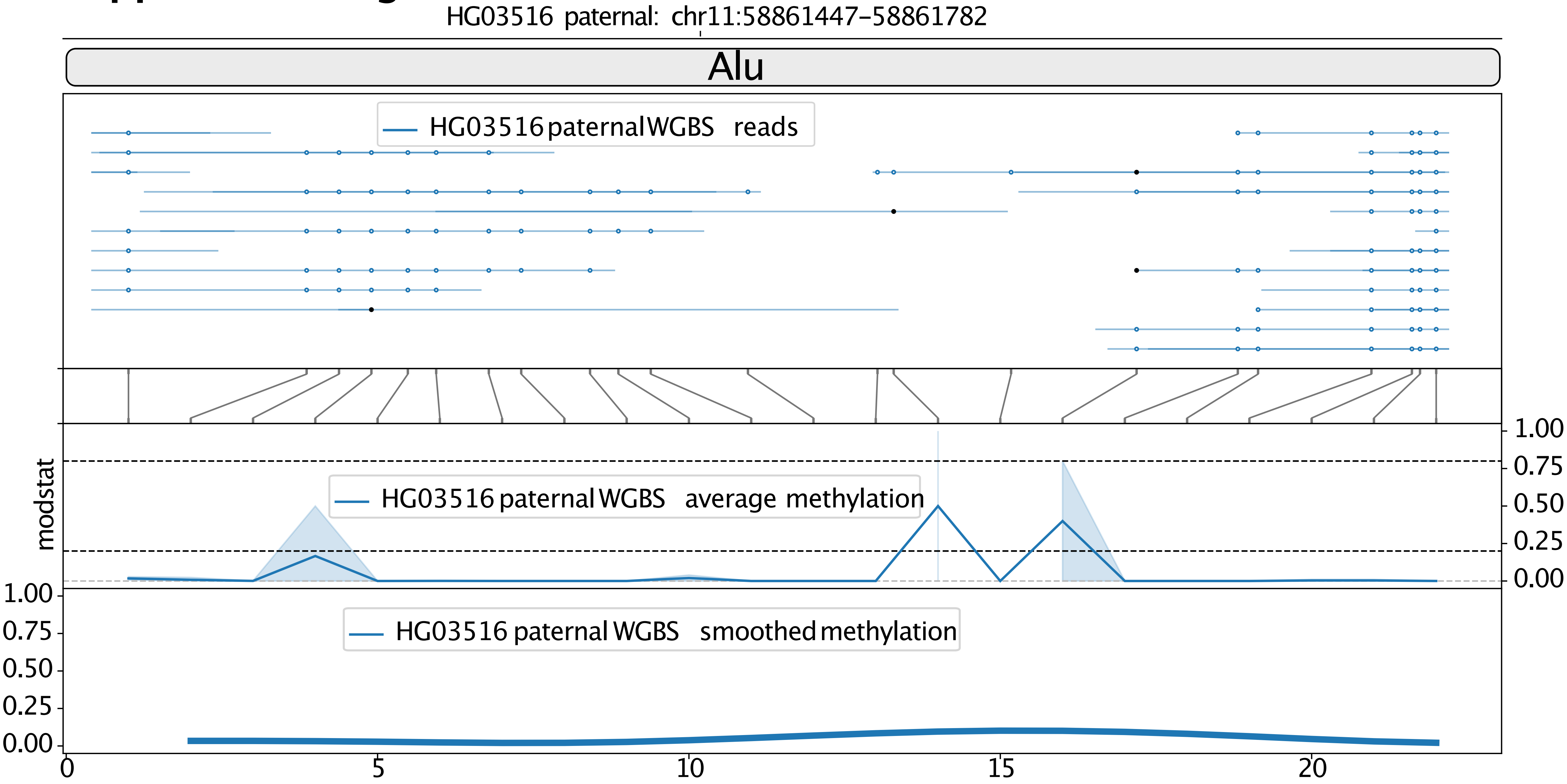

**Supplemental Fig. S3f**

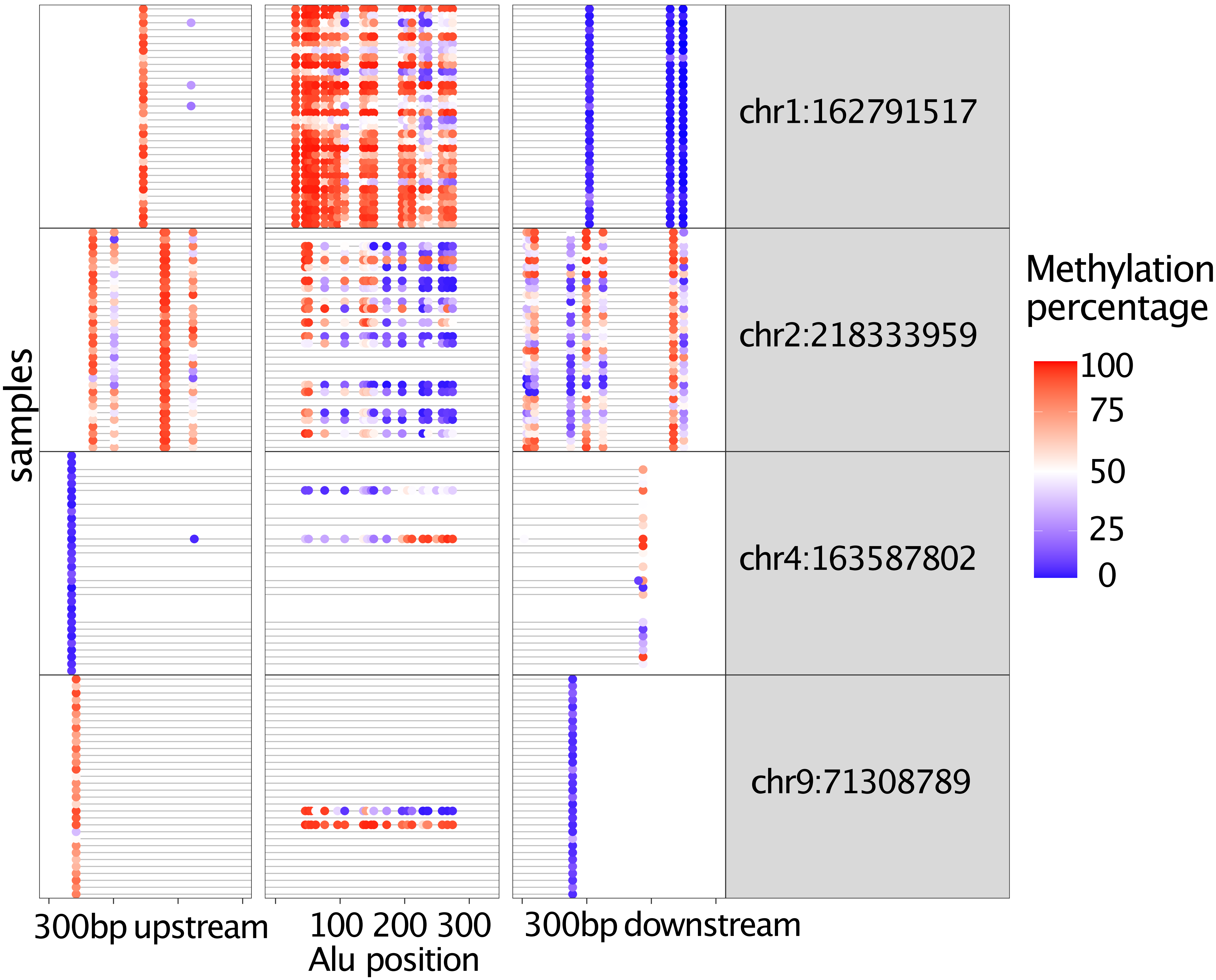

### Supplemental Fig. S3g

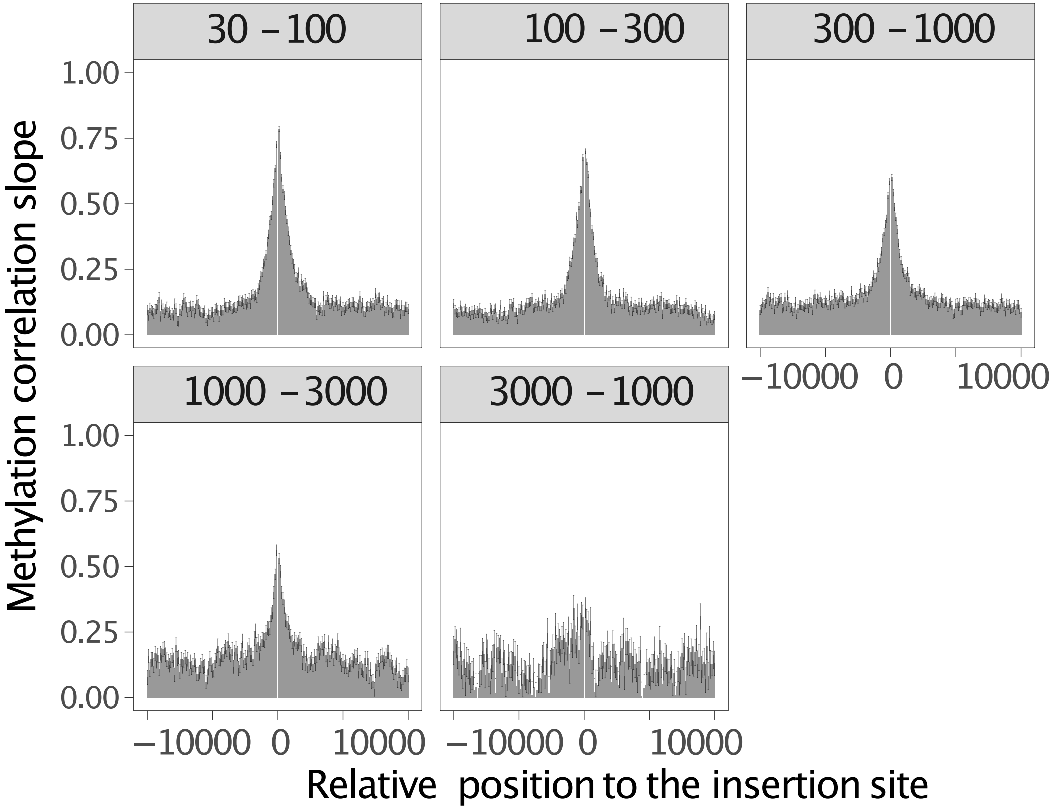
