## Supplementary figures and images for "Characterizing cytosine methylation of polymorphic human transposable element insertions using human pangenome resources"

### Supplemental Fig. S4

#### Supplemental Fig. S4a:

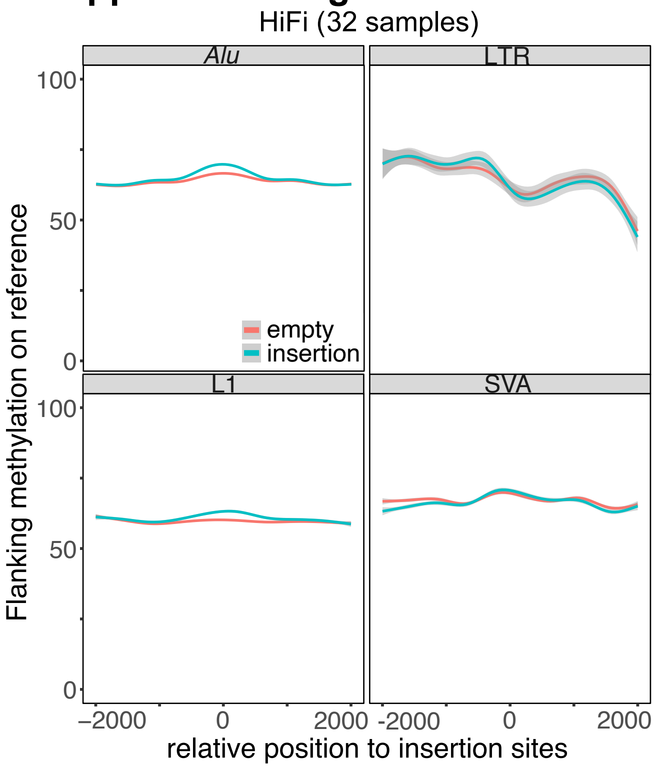


#### Supplemental Fig. S4b:

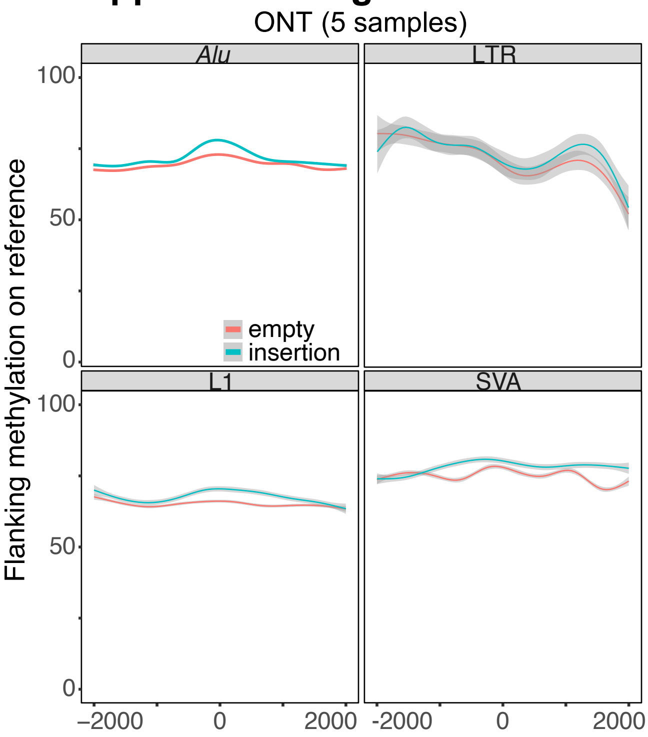


#### Supplemental Fig. S4c:

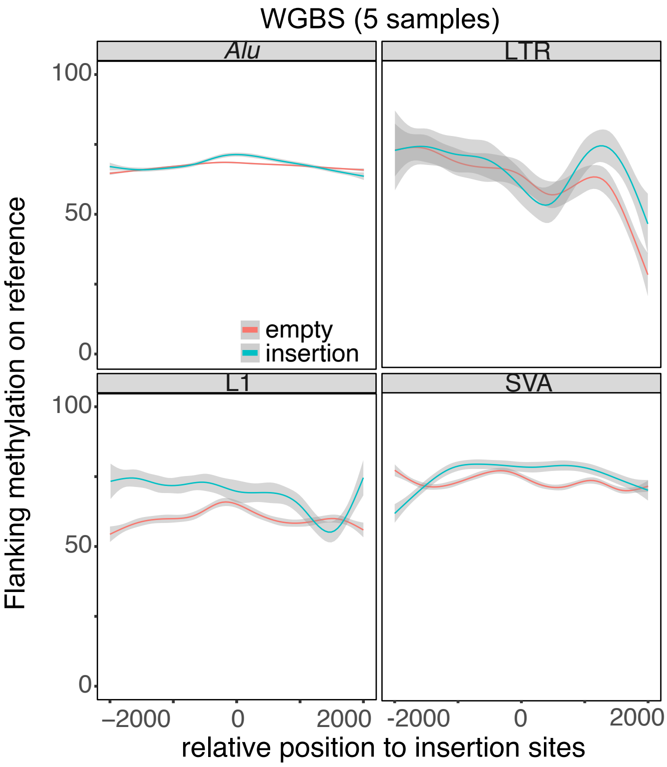

### Supplemental Fig. S6

**Supplemental Fig. S6:**

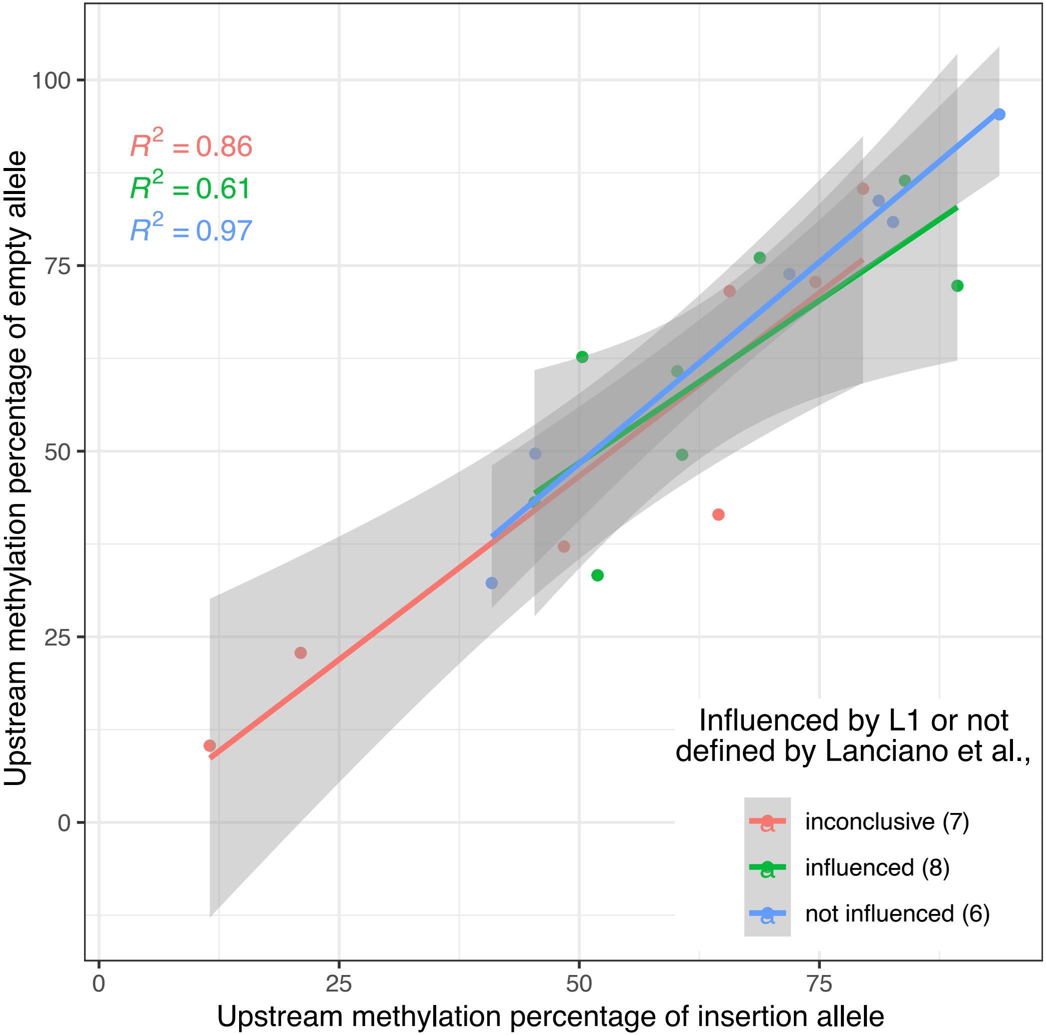
