## Supplemental Fig. S5 for "Characterizing cytosine methylation of polymorphic human transposable element insertions using human pangenome resources"

### Supplemental_Fig_S5a:

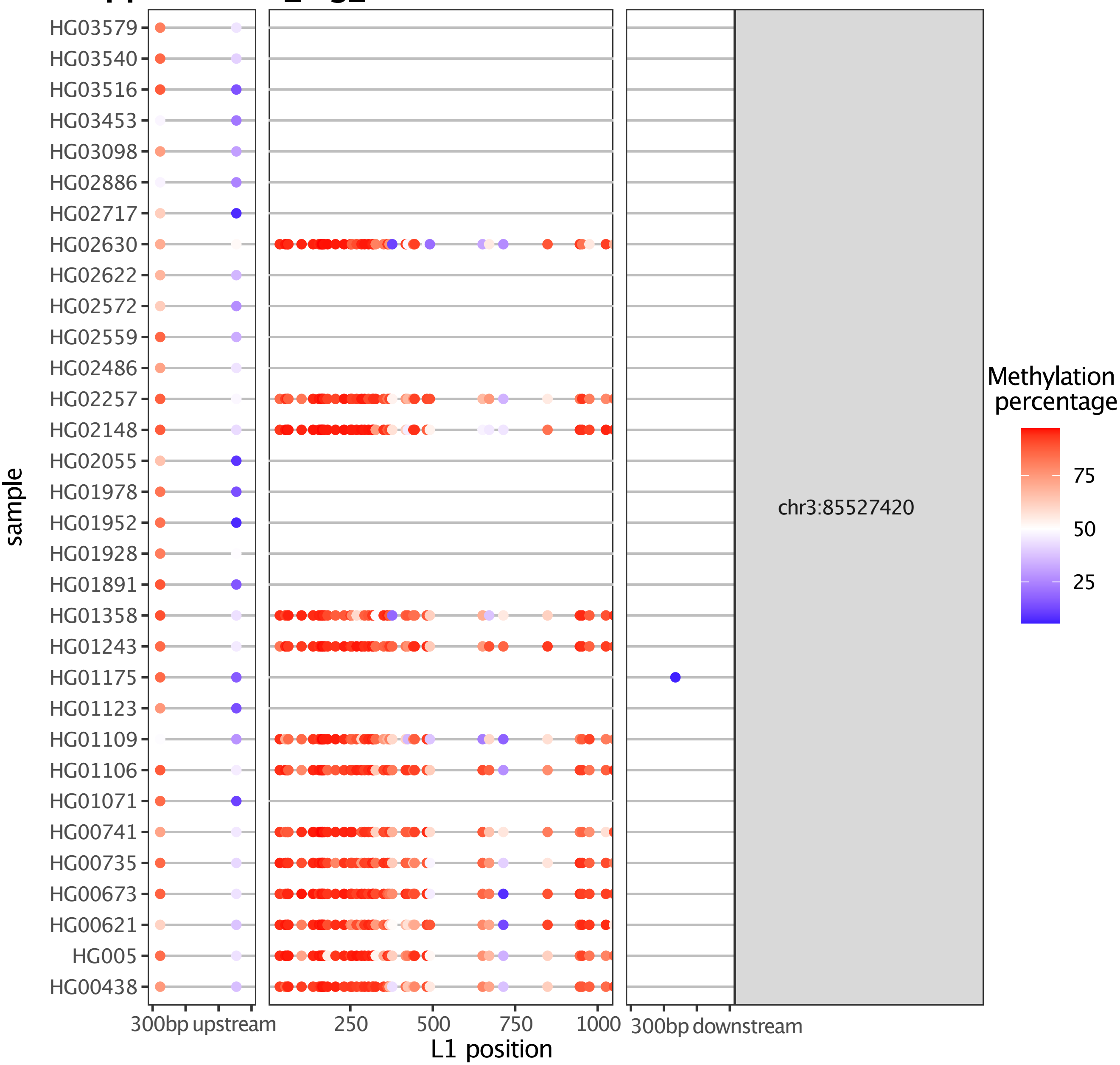

Supplemental_Fig_S5b:

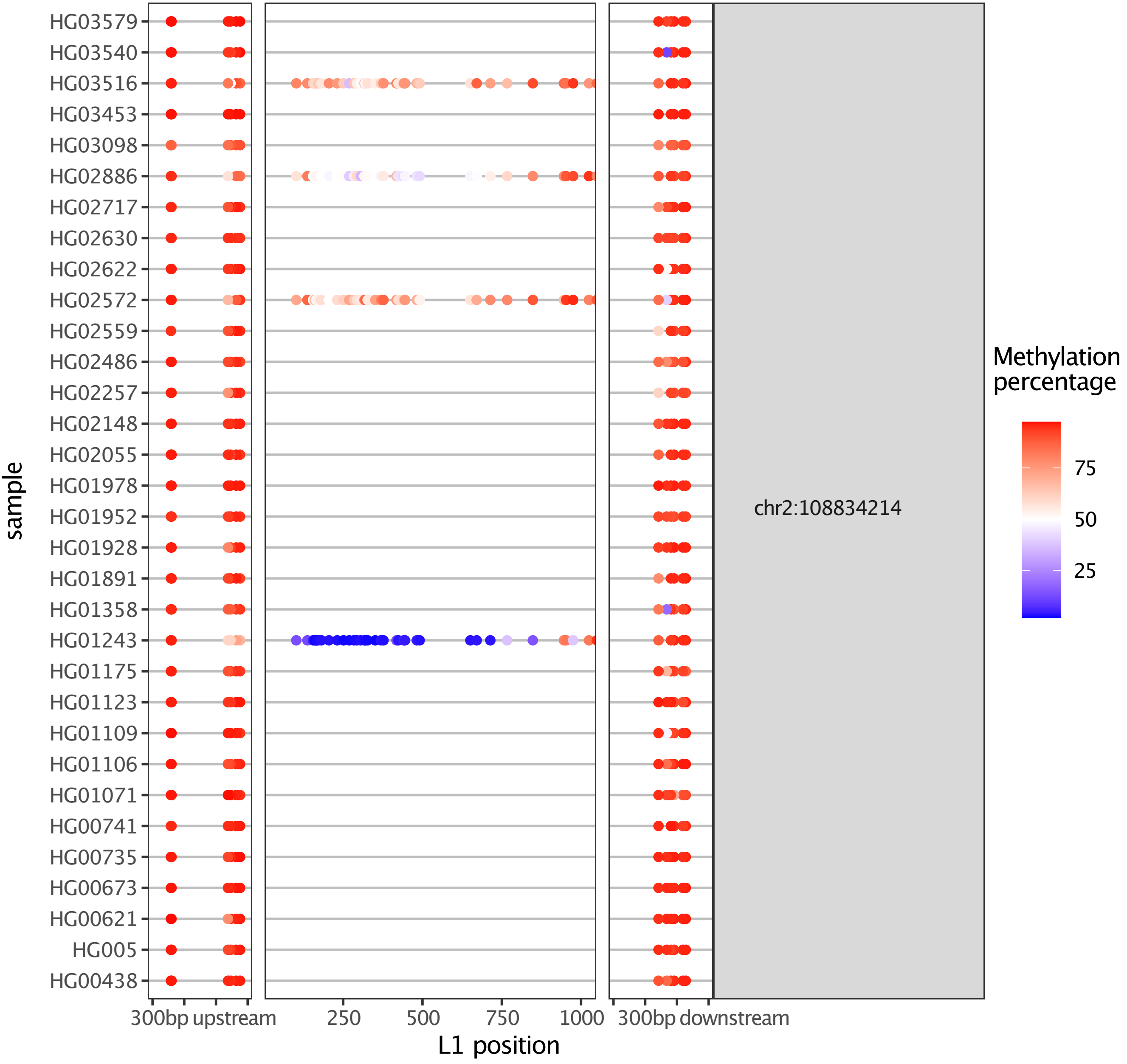

#### Supplemental_Fig_S5c:

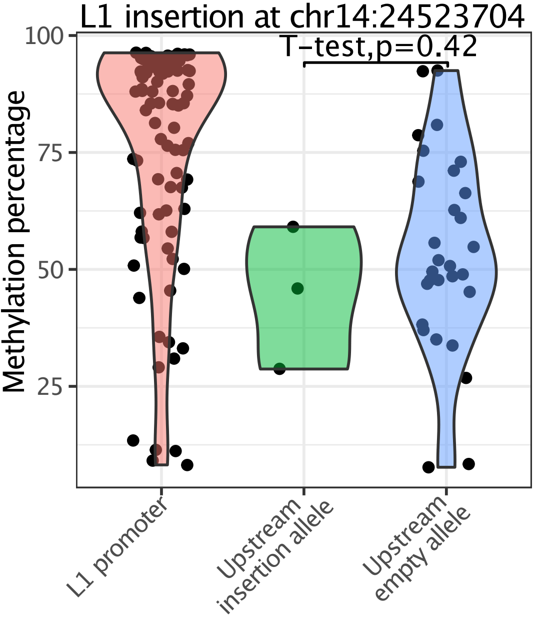

### Supplemental_Fig_S5d:

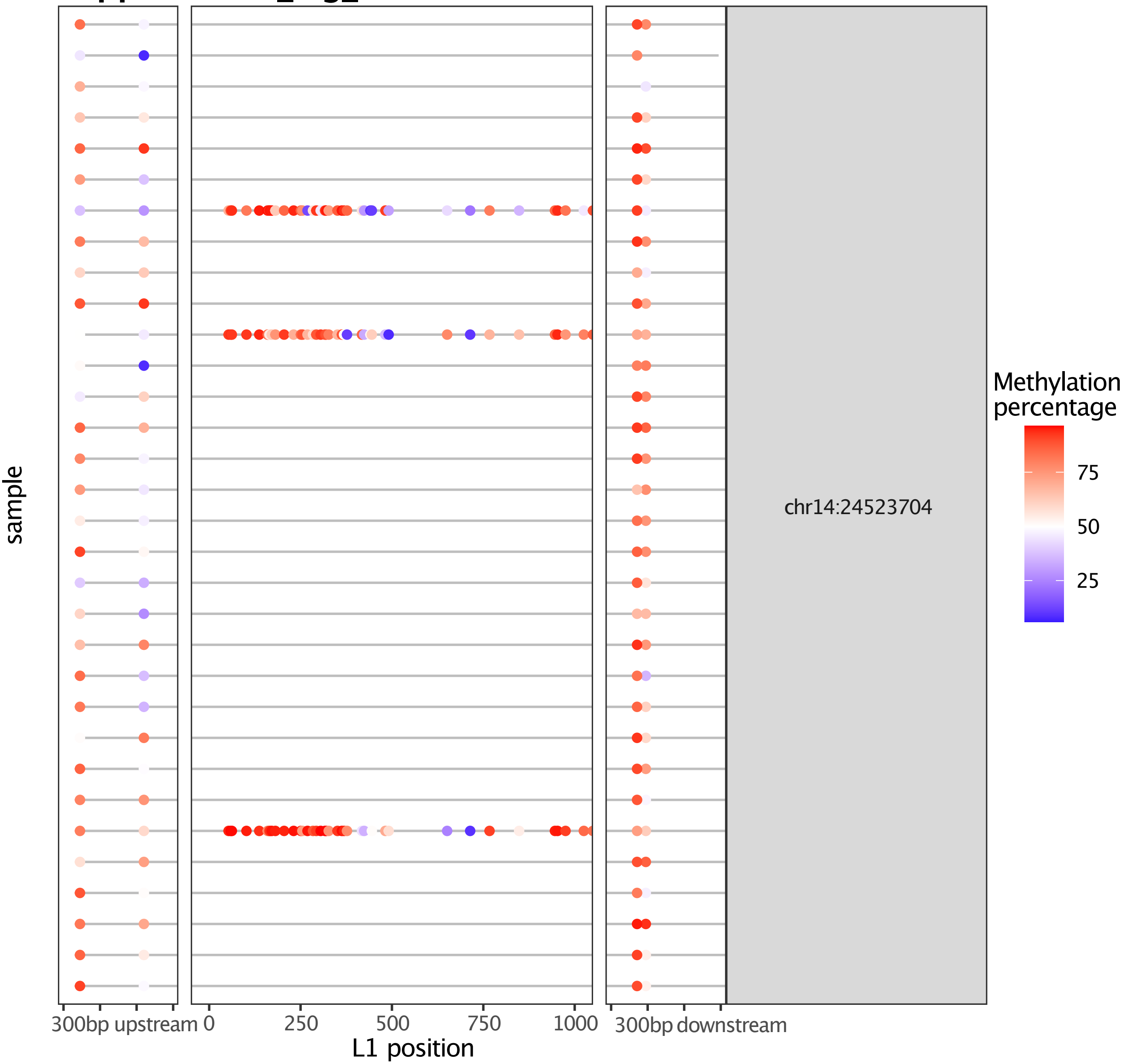

**Supplemental_Fig_S5e:**

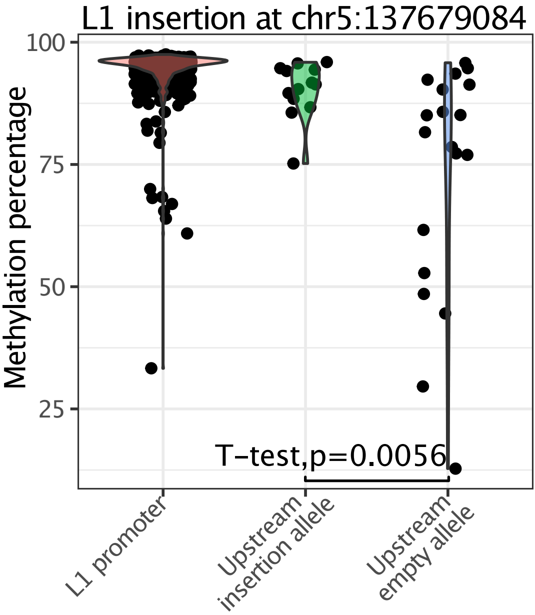

### Supplemental_Fig_S5f

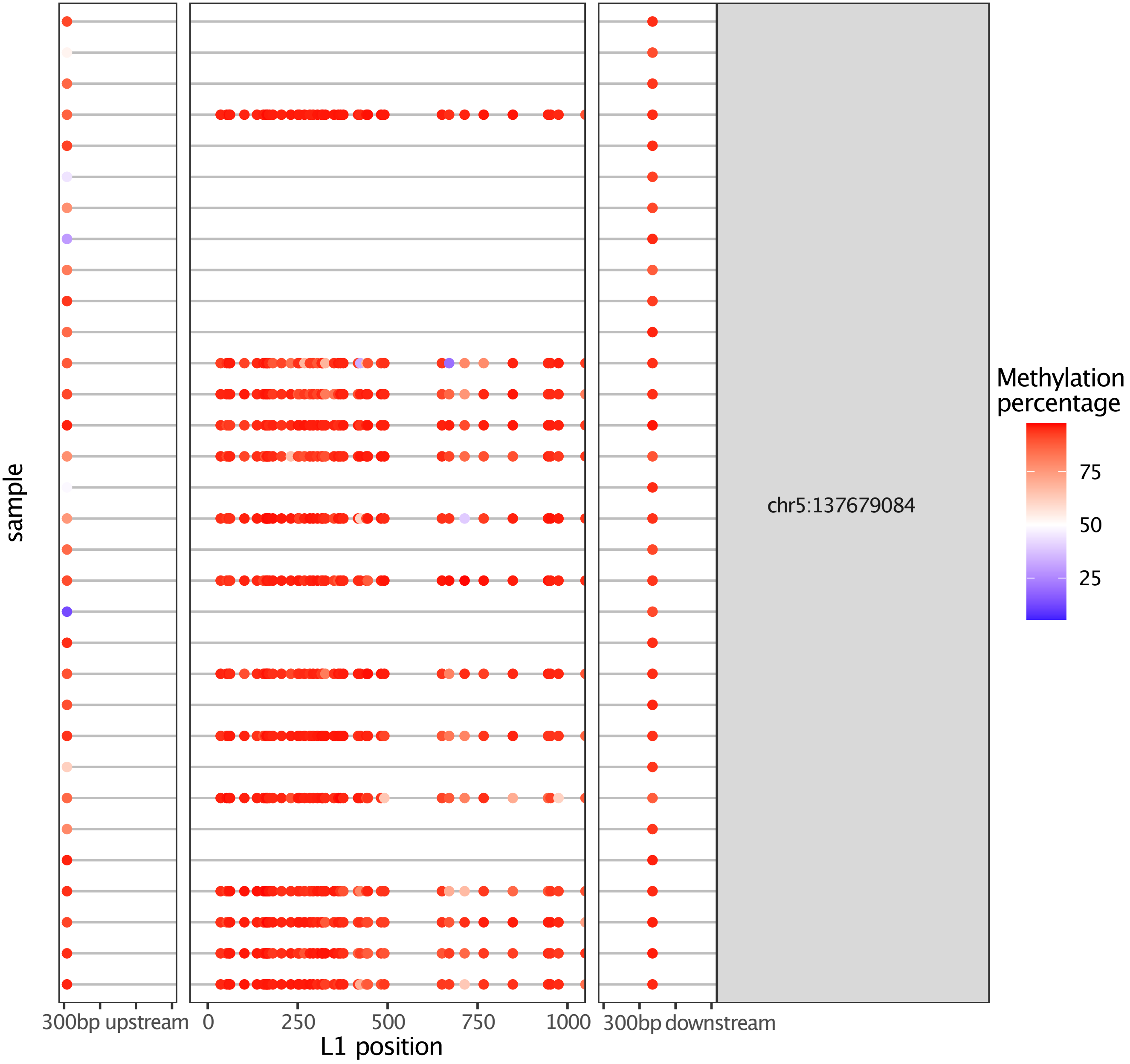

### Supplemental_Fig_S5g:

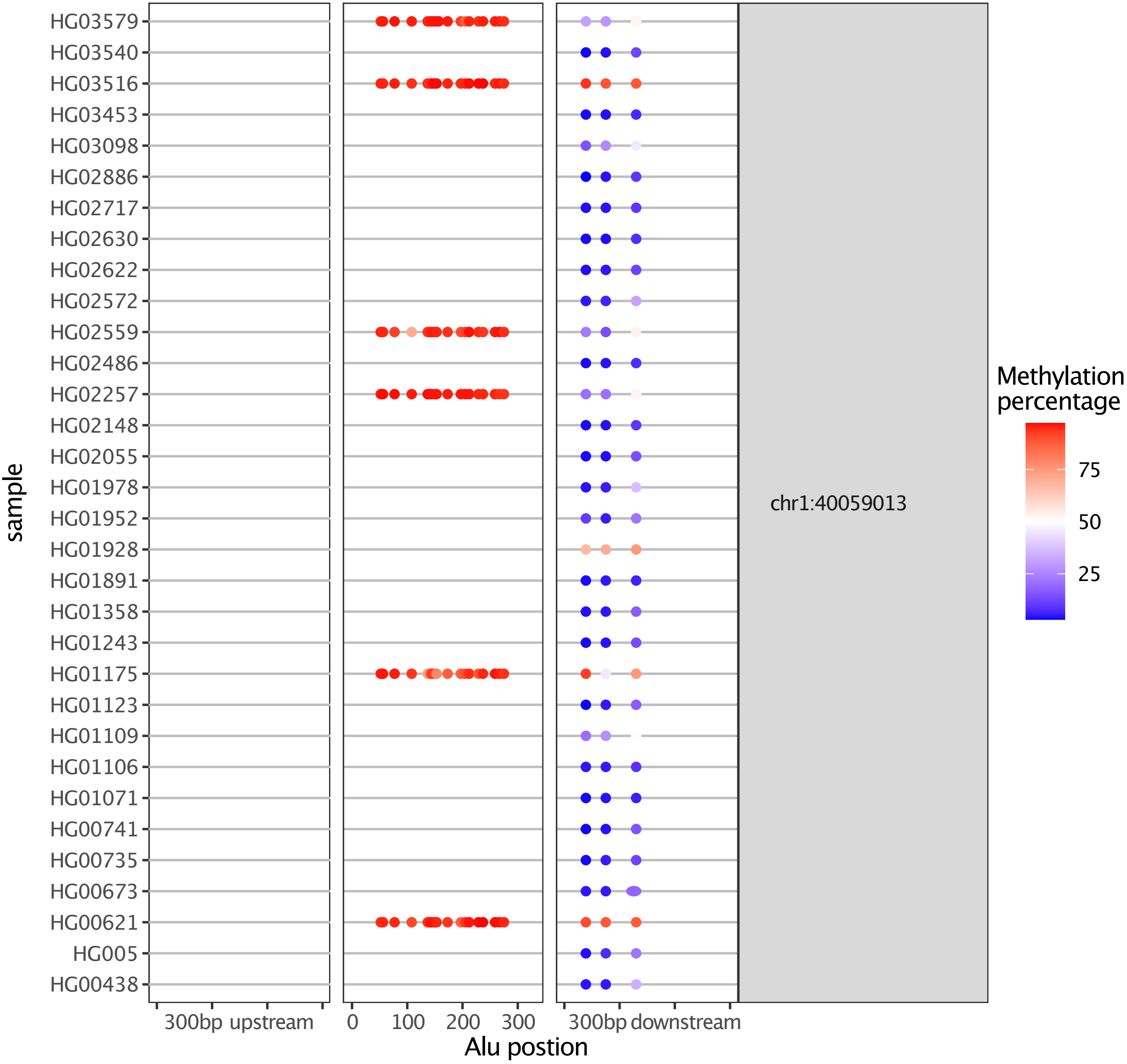

**Supplemental_Fig_S5h:**

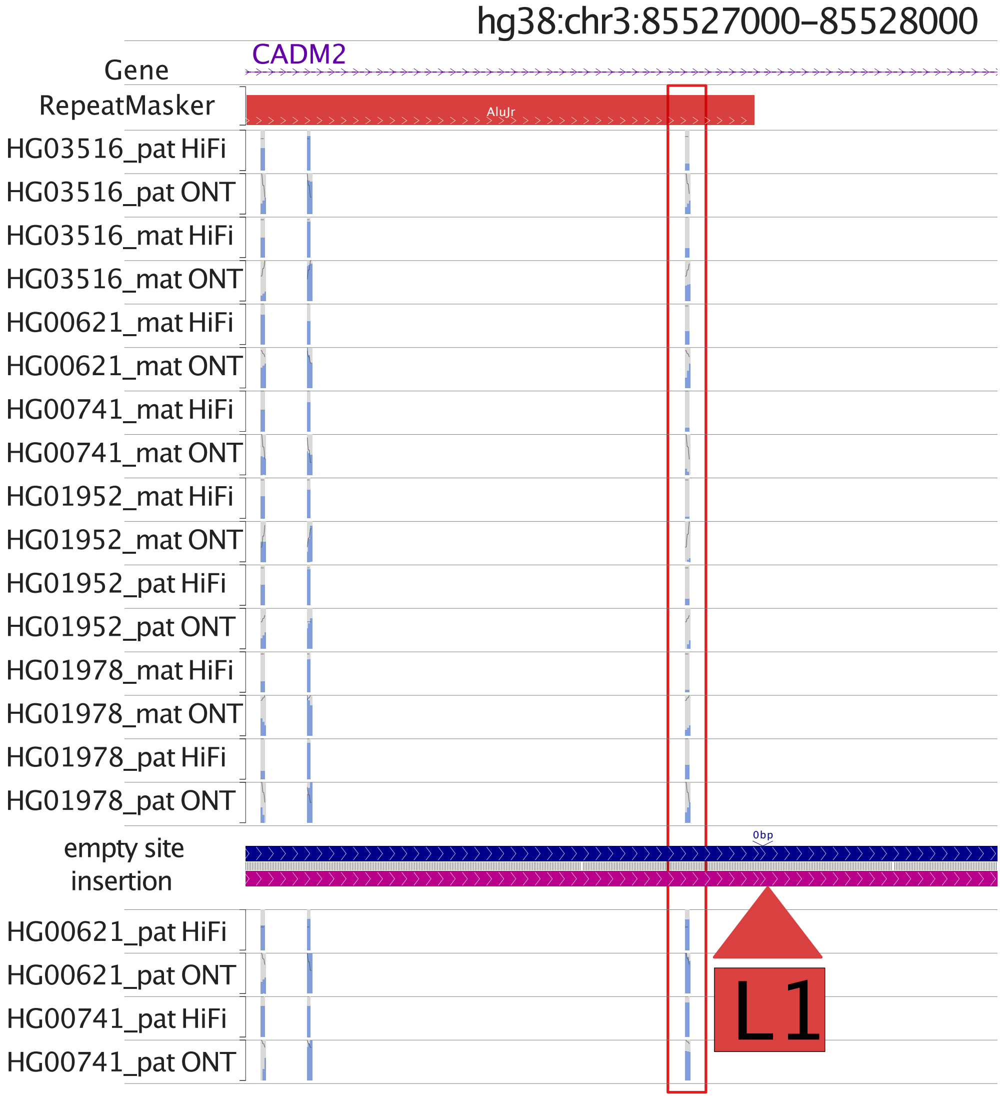
